# IGF2BP2 is Induced by Stress in the Heart and Mediates Dilated Cardiomyopathy

**DOI:** 10.1101/2022.11.03.515033

**Authors:** Miriam Krumbein, Froma Oberman, Yuval Cinnamon, Mordechai Golomb, Dalit May, Gilad Vainer, Vitali Belzer, Karen Meir, Irina Fridman, Johannes Haybaeck, Gerhard Poelzl, Izhak Kehat, Ronen Beeri, Sonja Kessler, Joel K. Yisraeli

**Author notes:** Both authors are senior authors (Sonja Kessler, Joel K. Yisraeli).

## Abstract

The IGF2BP family of RNA binding proteins consists of three paralogs that regulate intracellular RNA localization, RNA stability, and translational control. Although IGF2BP1 and 3 are oncofetal proteins, IGF2BP2 expression is maintained in many tissues, including the heart, into adulthood. Previous studies indicated that IGF2BP2 is upregulated in cardiomyocytes during cardiac stress and remodelling and returns to normal levels in recovering hearts. These results raise the possibility that IGF2BP2 might play an adaptive role during cardiac stress and recovery. Using a conditional, inducible transgenic mouse line, we found that enhanced expression of an IGF2BP2 transgene in newborn or adult hearts leads to dilated cardiomyopathy (DCM) and death within 3-4 weeks. Downregulation of the transgene after 2 weeks, however, rescues these mice, with complete recovery by 12 weeks. Hearts overexpressing IGF2BP2 downregulate sarcomeric and mitochondrial proteins and have fragmented mitochondria and elongated, thinner sarcomeres. Consistent with these results, IGF2BP2 is upregulated in patients with DCM or after myocardial infarction. These results suggest that IGF2BP2 may be an attractive target for therapeutic intervention in DCM.

## Introduction

IGF2BPs are a family of RNA binding proteins composed of three paralogs, IGF2BP1, IGF2BP2, and IGF2BP3, that are highly conserved from fish and amphibians to mammals (Yisraeli, 2005). Enhanced CLIP analysis of human embryonic stem cells indicates that the IGF2BP paralogs (particularly IGF2BP1 and IGF2BP2) overlap extensively in their RNA targets (Conway *et al*, 2016). Broadly expressed throughout embryonic development, IGF2BP1 and IGF2BP3 proteins are generally downregulated after birth, while IGF2BP2 expression is maintained in a number of adult tissues (Degrauwe *et al*, 2016). A wide variety of cancers show an upregulation of one or more of the IGF2BP proteins. The IGF2BP family of RNA binding proteins regulates RNA at many levels, including intracellular RNA localization, RNA stability, and translational control. The IGF2BPs are also reported to be readers of m^6^A RNA methylation, an epigenetic modification that can affect RNA stabilization and translation (Huang *et al*, 2018). Given the apparently large number of potential targets, it is not surprising that IGF2BP proteins have been implicated in many cellular functions, including cell migration, proliferation, axon turning and guidance, maintenance of epithelial structures, and metabolic regulation (Carmel *et al*, 2015; Dai *et al*, 2013; Janiszewska *et al*, 2012; Kalous *et al*, 2014; Laggai *et al*, 2014).

Cardiac disease, one of the leading causes of death worldwide, affects millions of people yearly. RNA binding proteins (RBPs) have come under increased scrutiny for their role in cardiac health and disease (Akerberg *et al*, 2019; Gao & Wang, 2020; Liao *et al*, 2016). Recently, a network of RBPs has been shown to associate with α−actinin in iPS-induced cardiomyocytes, helping to transport mitochondrial RNAs to the sarcomere and aiding in metabolic adaptations to stress (Ladha *et al*, 2021). Among these RBPs is IGF2BP2.

We report here that IGF2BP2 is normally expressed at low levels in postnatal and adult hearts, but that pressure overload or ischemia, both of which cause hypertrophy, lead to an increase of IGF2BP2 levels. Furthermore, activating IGF2BP2 expression in transgenic mice results in dilated cardiomyopathy within 2 to 3 weeks. The initial response to IGF2BP2 activation leads to heart failure or cardiac dysfunction and precocious death within 3 to 4 weeks after induction of IGF2BP2 expression. In both newborn and adult mice, induced expression leads to a progressive loss of heart function, but this process is reversible if exogenous IGF2BP2 expression is turned off in time. Adult hearts in which IGF2BP2 expression is activated for two weeks show a striking decrease in mitochondrial and sarcomeric proteins, as well as a shift in the expression of proteins associated with mitochondrial to those associated with non-mitochondrial based metabolism. Analysis of EM sections of these hearts reveals a preponderance of fragmented mitochondria and elongated, thinner sarcomeres, validating these findings. Consistent with the mouse data, elevated levels of IGF2BP2 are observed in human patients with dilated cardiomyopathies or after myocardial infarctions. These results suggest a model in which IGF2BP2 is upregulated upon various cardiac stresses and induces remodelling as part of an adaptive response. This response, however, comes at the price of impaired mitochondria and reduced sarcomere organization that eventually leads to heart failure (cardiac dysfunction) if IGF2BP2 levels are not reduced.

## Results

### IGF2BP2 is upregulated as a result of cardiac stress

To better understand the role of IGF2BPs in the heart, we characterized the expression of all three IGF2BP paralogs in maturing hearts at both the RNA and protein level. As expected for oncofetal proteins, IGF2BP1 and IGF2BP3 RNA and protein levels decline rapidly after birth, with essentially baseline levels observed by 3 weeks of age. IGF2BP2 RNA expression, however, is maintained even in the adult heart, with a low but detectable level of protein still observed at 6 weeks (Fig. 1A, B).

**Figure 1.**
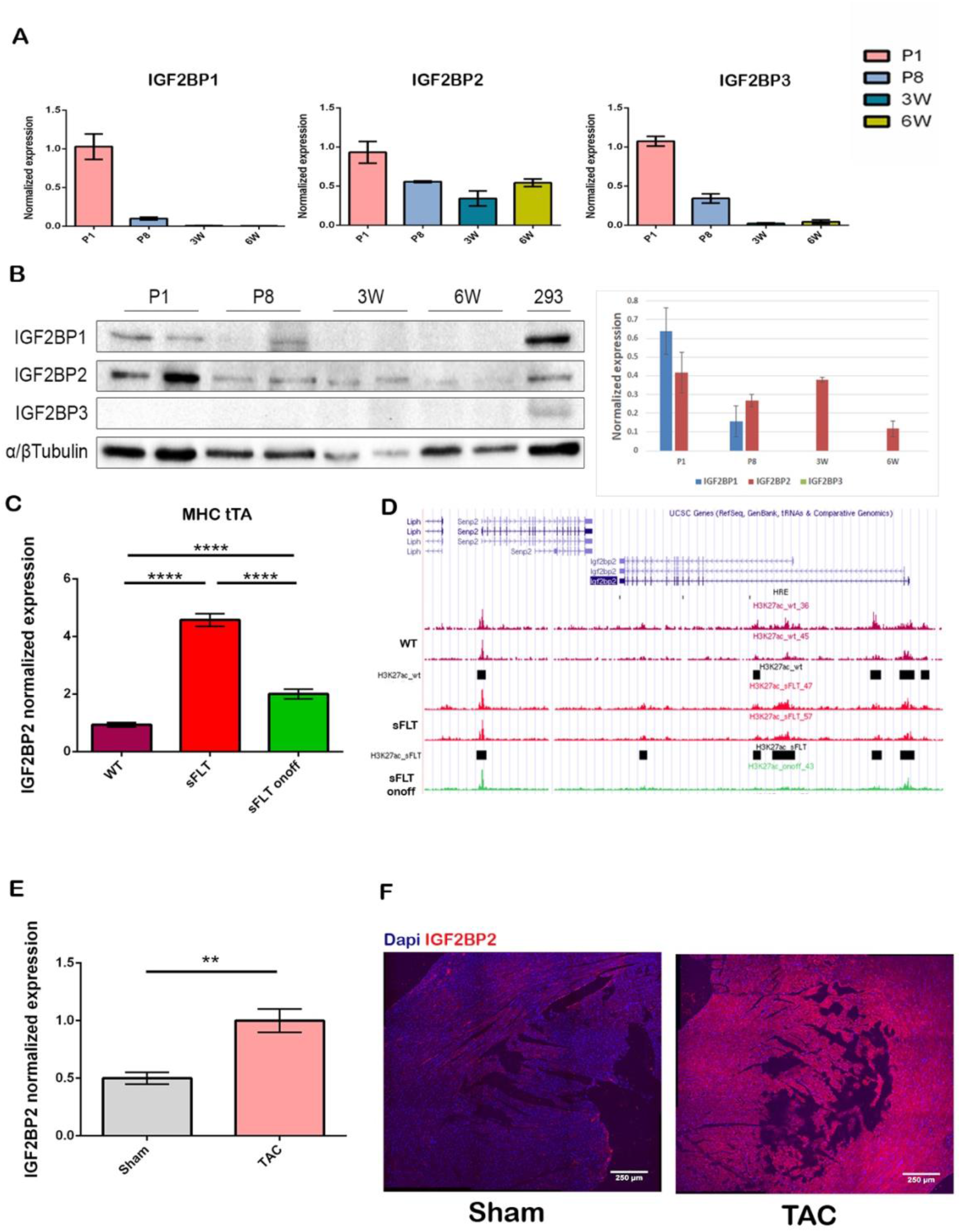
Expression of IGF2BP paralogs and cardiac stress models. (A) Relative IGF2BP2 RNA expression was measured by RT-PCR in normal hearts at post-natal day 1 (P1), postnatal day 8 (P8), 3 weeks (3W) and 6 weeks (6W) of age. Each bar represents three biological repeats ± SEM). (B) Western blot of IGF2BPs at the indicated time points. α/β Tubulin was used as a loading control. On the right, quantification of western results. Two biological repeats are shown for each time point ± SEM. (C) IGF2BP2 RNA expression in hearts from wildtype (WT), sFlt upregulated (sFLT), and sFlt upregulated and then downregulated (sFLT onoff) was compared on a gene array (*p*<0.0001). (D) CHIP analysis of H3K27ac - enhancer marks in the genomic region of IGF2BP2 in hearts isolated from WT, sFLT upregulated, and sFLT upregulated and then downregulated mice. Black boxes represent significant peaks. (E) IGF2BP2 mRNA expression from sham or TAC mice was assayed by RT-PCR (n=3; *p*=0.0015). (F) IGF2BP2 expression was detected by immunofluorescence (red staining) on heart sections from sham-operated or TAC mice. Nuclei are stained in blue (DAPI). Scale bar is 250µm. Statistical significance was determined by two tailed Student’s t-test. ** *p*<0.01, \*\*\*\**p*<0.0001. *A. U*., arbitrary units.

Cardiomyopathy, which leads to heart failure and death, can be induced by both ischemic and non-ischemic stress to the heart. To investigate whether IGF2BP proteins might be activated during cardiac stress, we initially searched available RNA expression data from various cardiac stress models for expression of any of the IGF2BP paralog RNAs. The only paralog that was significantly upregulated in several mouse models of induced cardiac stress was IGF2BP2. Temporary ligation of the left anterior descending (LAD) coronary artery creates ischemia in the left ventricle, and this technique has been used as a model to induce myocardial infarction (Kolk *et al*, 2009). IGF2BP2 RNA is upregulated specifically in LAD ligated, but not sham-operated, mice at 24 and 48 hours post occlusion, and only distal to the ligation (Fig. S1A) (Tarnavski *et al*, 2004). Significantly, IGF2BP2 expression returns to normal levels by 8 weeks, concomitant with recovery. A similar correlation between elevated IGF2BP2 expression and dilated cardiomyopathy and heart failure, followed by reduced IGF2BP2 expression upon resolution of heart failure, is observed in MerCreMer mice (Koitabashi *et al*, 2009) (Fig. S1B). IGF2BP2 is also activated by isoproterenol (Fig. S1C (Galindo *et al*, 2009)) and phospholamban-induced dilated cardiomyopathies (Burke *et al*, 2016b).

We examined the response of IGF2BP2 to cardiac stress in two mouse models. In the first model, inducible expression of a soluble VEGF (sFlt) receptor in the heart of juvenile mouse pups causes chronic hypoxia, cardiac remodelling and hibernation of cardiomyocytes (May *et al*, 2008). When we analyzed the data from these mice, we found that IGF2BP2 RNA expression is upregulated over 4-fold as a result of this induced ischemia. Silencing of the sFlt receptor, which rescues the phenotype, concomitantly downregulates IGF2BP2 expression (Fig.1C). Furthermore, ChIP analysis of the IGF2BP2 gene reveals that H3K27 acetylation, a marker of transcriptional activation, varies with the activation and silencing of the sFlt receptor and mirrors mRNA expression levels of IGF2BP2 (Fig.1D). In a second model (deAlmeida *et al*, 2010), transverse aortic constriction (TAC) generates pressure overload in the heart, a procedure that leads to cardiac hypertrophy and heart failure. RT-PCR performed on RNA isolated from these hearts revealed an upregulation of IGF2BP2 RNA as compared to the sham-treated hearts (Fig.1E). Upregulation of IGF2BP2 protein was also observed by immunofluorescent staining of sections from TAC versus sham-treated mice (Fig.1F). Taken together, these experiments indicate that various types of stress that lead to cardiomyopathies and heart failure upregulate IGF2BP2 in the heart.

### IGF2BP2 upregulation in the heart leads to cardiac remodelling, fibrosis, heart failure and death

Cardiac stress can have profound and pleiotropic effects on heart physiology. To test directly the effect of IGF2BP2 upregulation in the heart, we used a transgenic system that enabled us to express human IGF2BP2 (hIGF2BP2) specifically in cardiac tissue in a conditional and reversible manner, in the absence of any induced stress. Transgenic mice expressing a tetracycline-regulated transactivator protein (tTA) from a heart-specific myosin heavy chain (MHC) α promoter were mated with mice harboring a transgene encoding an hIGF2BP2 protein under control of the tetracycline response element upstream of a minimal cytomegalovirus (TRE-CMVmin) promotor (Fig. 2A) (see Materials and Methods). In offspring possessing both transgenes, cardiac-specific expression of hIGF2BP2 can be regulated by tetracycline through the addition of, or withdrawal from, the drinking water (off or on, respectively). Transgene expression in cardiac tissue from these mice was confirmed by histological staining (Fig. S2A), and the level of expression of the hIGF2BP2 transgene was approximately 2.5 fold above that of the endogenous mouse IGF2BP2, as determined by RT-PCR using RNA isolated from primary cardiomyocytes grown from mice induced for 5 days (Fig. S2B).

**Figure 2.**
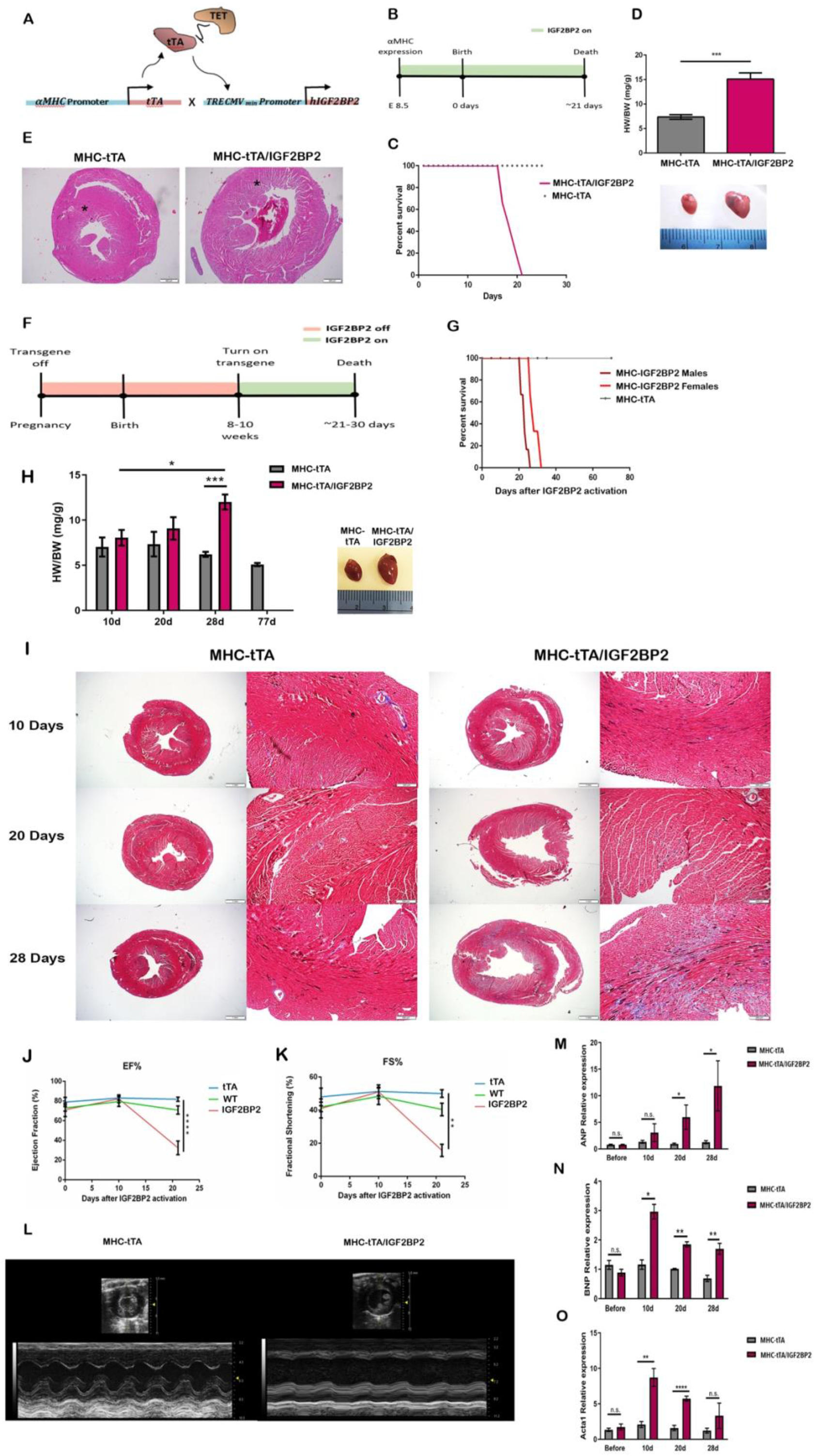
Activation of the conditional IGF2BP2 transgene in mouse hearts. (A) Schematic representation of the transgenic mouse cross. (B) Timeline of transgene expression for the hearts shown in C-E. (C) Kaplan–Meier survival curve of the transgenic mice. n= 24 (MHC-tTA/IGF2BP2=6, WT=6, MHC-tTA=6, TRE-IGF2BP2=6). (D) Bar graph of the ratio of heart weight to body weight in MHC-tTA mice and MHC-tTA/IGF2BP2 mice. Below the graph are representative pictures of an MHC-tTA (left) heart and an MHC-tTA/IGF2BP2 heart (right) ± SEM (MHC-tTA/IGF2BP2=4, WT=3, MHC-tTA=2, TRE-IGF2BP2=3) (*p*=0.0001). (E) H&E staining of a cross section of an MHC-tTA (left) and MHC-tTA/IGF2BP2 heart (right). Scale bar is 500µm. Asterisks indicate the sub-endocardium. (F) Timeline of transgene expression for the hearts used in the experiments described in G-O. (G) Kaplan–Meier survival curve of adult transgenic mice. (MHC-tTA/IGF2BP2, males n=6, females n=6; MHC-tTA, males n=6, females n=5). (H) Bar graph of the ratio of heart weight to body weight (HW/BW) (mg/g) ± SEM in adult MHC-tTA mice and MHC-tTA/IGF2BP2 mice at different time points after transgene induction (10d: MHC-tTA, n=3; MHC-tTA/IGF2BP2, n=5; 20d: MHC-tTA, n=3; MHC-tTA/IGF2BP2, n=4; 28d: MHC-tTA, n=5; MHC-tTA/IGF2BP2, n=3) (10d-20d, p=0.0233, 28d, *p*=0.0002). A representative picture of an MHC-tTA heart (left) next to an MHC-tTA/IGF2BP2 heart (right) at end point. (I) Trichrome staining of MHC-tTA (left) and MHC-tTA/IGF2BP2 (right) hearts at the indicated time points. Scale bar is 1mm and 100µm in the higher magnification (right panels). (J-K) Echocardiograms were performed on WT, MHC-tTA, and MHC-tTA/IGF2BP2 mice before, 10 days post, and 21 days post activation of the transgene (n=6 in each group). Analyses were of (J) Ejection fraction percentage and (K) Fractional shortening percentage. Statistical significance was determined by one-way analysis of variance (ANOVA) followed by Dunnett’s post hoc test. (L) Representative pictures of the echocardiogram at 21 days post activation. (M-O) RNA was extracted from MHC-tTA and MHC-tTA/IGF2BP2 hearts at the indicated time points and analysed by RT-PCR for expression of ANP (20d, *p*=0.0462; 28d, *p*=0.0226), BNP (10d, *p*=0.0136; 20d, *p*=0.0034; 28d, *p*=0.0039), and ACTA1 (10d, *p*=0.0076; 20d, *p*<0.0001). Values show the results from 2 biological repeats ± SEM. Statistical significance in the bar graphs was determined by two tailed Student’s t-test. * *p*<0.05, ** *p*<0.01, \*\*\**p*<0.001, \*\*\*\**p*<0.0001.

When tetracycline is not administered to pregnant mice containing both transgenes, hIGF2BP2 expression is activated at the time that the MHCα gene is first turned on, around day E8.5 (Ishikawa *et al*, 2003) of development (Fig.2B). Pups appear normal at birth but all of them die very early, even before weaning, beginning at 21 days (Fig.2C). When sacrificed at twelve days of age, these mice have enlarged and remodelled hearts (Fig.2D). In histological sections stained for H&E, it is clearly apparent that the hIGF2BP2-expressing hearts are much larger, and that their endo and sub-endocardium are very sparse compared to those of control hearts (Fig.2E).

To determine whether upregulation of IGF2BP2 can affect not only perinatal but also adult hearts, mice were maintained on tetracycline for 8-10 weeks (IGF2BP2 off) before the antibiotic was removed from their drinking water (Fig.2F). Induction of hIGF2BP2 at this point also leads to heart failure and death in all the mice, although the process takes slightly longer in females as opposed to males (26-32 days as opposed to 21-26 days, respectively). As observed in the perinatally induced mice, the hearts were enlarged and remodelled, and these modifications progressed gradually from 10 to 28 days post-hIGF2BP2 induction (Fig.2H). In histological slides prepared from these hearts at the different time points, the changes observed at 10 days post-induction are very subtle, with a slight disarray and mismatch of cell size of the cardiomyocytes, mainly in the papillary muscle and the sub-endocardium. With time, these differences become more pronounced, and fibrosis increases, also localizing to the sub-endocardium close to the lumen of the left ventricle (Fig.2I and Supp.2C).

Heart function was monitored in these mice by echocardiography (Fig.2J-L). Prior to induction of the IGF2BP2 gene, the hearts functioned normally in each of the three cohorts (MHC-tTA/IGF2BP2, MHC-tTA and WT), indicating that there was no leakiness of hIGF2BP2 expression. At 10 days post-induction, although the cardiac stress/DCM markers ANP, BNP, and alpha actin 1 are already upregulated at the transcriptional level (Fig.2 M-O), the heart still appears to function normally, with no apparent differences observed in ejection fraction or fractional shortening. At 21 days post induction, however, mice expressing hIGF2BP2 exhibit a clear DCM phenotype (McNally & Mestroni, 2017), with ejection fraction under 40%, fractional shortening under 20%, LV end diastolic diameter enlarged by more than 120% (Fig.2J-L), and stress markers at elevated levels. These results argue that upregulation of IGF2BP2 expression rapidly induces stress in cardiomyocytes and is sufficient to cause DCM progressing to heart failure and death in both perinatal and adult hearts within 3-4 weeks.

### IGF2BP2-induced cardiomyopathy is reversible

In both the LAD occlusion model of MI (Fig. S1A) and the MerCreMer model (Fig. S1B), IGF2BP2 appears to return to nearly normal levels several weeks after the initial stress, and this drop correlates with rescue of the cardiac myopathy phenotype. These findings prompted us to test whether cessation of hIGF2BP2 expression after induction of cardiac remodelling could rescue mice from progressing to complete heart failure. We employed two groups of transgenic mice. In the first group, hIGF2BP2 expression was induced for 2 weeks in 11 adult mice (8-10 weeks old) and then switched off (Fig.3A). No loss of heart function was observed (Fig.3B, C), and there was a 100% survival rate. This was in strikingly contrast to the mice in which hIGF2BP2 was maintained beyond 2 weeks and the survived rate was 0% after 32 days (Fig. 2). Three of the mice in the group that survived were sacrificed 4 weeks later, and their hearts were found to be slightly enlarged, indicating that some remodelling had occurred as a result of the 2 week induction (Fig.3D). In these hearts, wheat germ agglutinin staining of the endocardium adjacent to the papillary muscle confirmed that the cardiomyocytes were hypertrophic (Fig. 3E). The 8 remaining mice survived until they were sacrificed, 12 weeks after the gene had been turned off. Heart function was not affected throughout the entire post-IGF2BP2 expression period (Fig.3B, C), and their hearts returned to normal size (Fig.3F). These findings indicate that the effects of IGF2BP2 expression on cardiac remodelling and function are potentially reversible.

**Figure 3.**
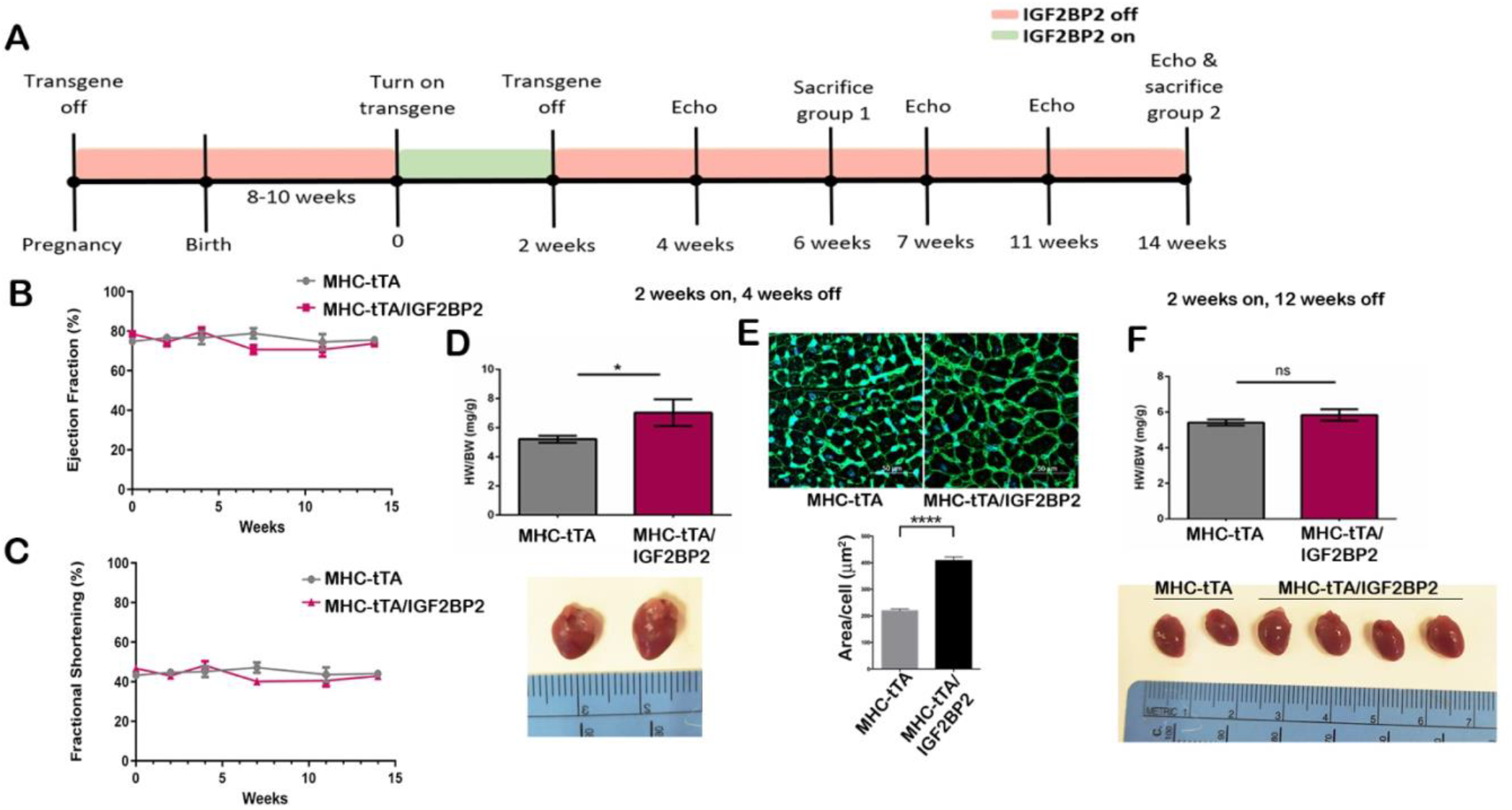
Rescue experiment with full recovery. (A) Timeline of experimental design for the experiments described in B-F: transgene on for 2 weeks and off for 12 weeks in adult mice. (B-C) Echocardiograms were performed on all the mice over the course of the experiment and analyzed for Ejection Fraction percentage (B) and Fractional Shortening percentage (C). (D) Bar graph of the ratio of heart weight to body weight (HW/BW) (mg/g) ± SEM in mice after 2 weeks of induction and 4 weeks after turning off the transgene. Below the graph are representative pictures of an MHC-tTA heart (left) next to an MHC-tTA/IGF2BP2 heart (right) (*p* =0.0299) (MHC-tTA, n=9; MHC-tTA/IGF2BP2, n=5). (E) Wheat germ agglutinin staining of endocardial tissue adjacent to the papillary muscle from the hearts described in (D). Quantification of the area of 150 cardiomyocytes from each cohort is shown below. (F) Bar graph of the ratio of heart weight to body weight (HW/BW) (mg/g) ± SEM in mice after 2 weeks of induction and 12 weeks after turning off the transgene. Below the graph are representative pictures of two MHC-tTA (left) and four MHC -tTA/IGF2BP2 hearts (right) (MHC-tTA n=5, MHC-tTA/IGF2BP2, n=8). Statistical significance was determined by two tailed Student’s t-test. * p<0.05, **** p<0.0001.

A second group of 5 mice were induced for hIGF2BP2 expression for 16 days (Fig.4A). As in the first group, at the end of the induction there was no loss of heart function (Fig.4B,C). Nevertheless, the outcome in this group varied significantly. Two mice (I and II) died within two weeks of turning off hIGF2BP2. The heart from mouse II had extensive dilation of both ventricles and loss of muscle wall thickness (Fig.4F). Although the remaining three survived until the end of the experiment, at 12 weeks post-hIGF2BP2 expression, one of the mice (mouse III) had severely impaired heart function (ejection fraction and fractional shortening; Fig. 4B, C) and a significantly large heart with extensive fibrosis throughout all of the left ventricle wall (Fig. 4E, F). The remaining two mice (IV and V) displayed some impairment in heart function that almost reverted back to normal by the end of the experiment, and their hearts were of normal size and with minimal fibrosis (Fig. 4D, F and data not shown). The variability among this set of mice suggests that at some time slightly after 14 days of hIGF2BP2 expression, remodelling of the heart progresses beyond a point of no return when the damage to the heart cannot be overcome, even if the gene is turned off.

**Figure 4.**
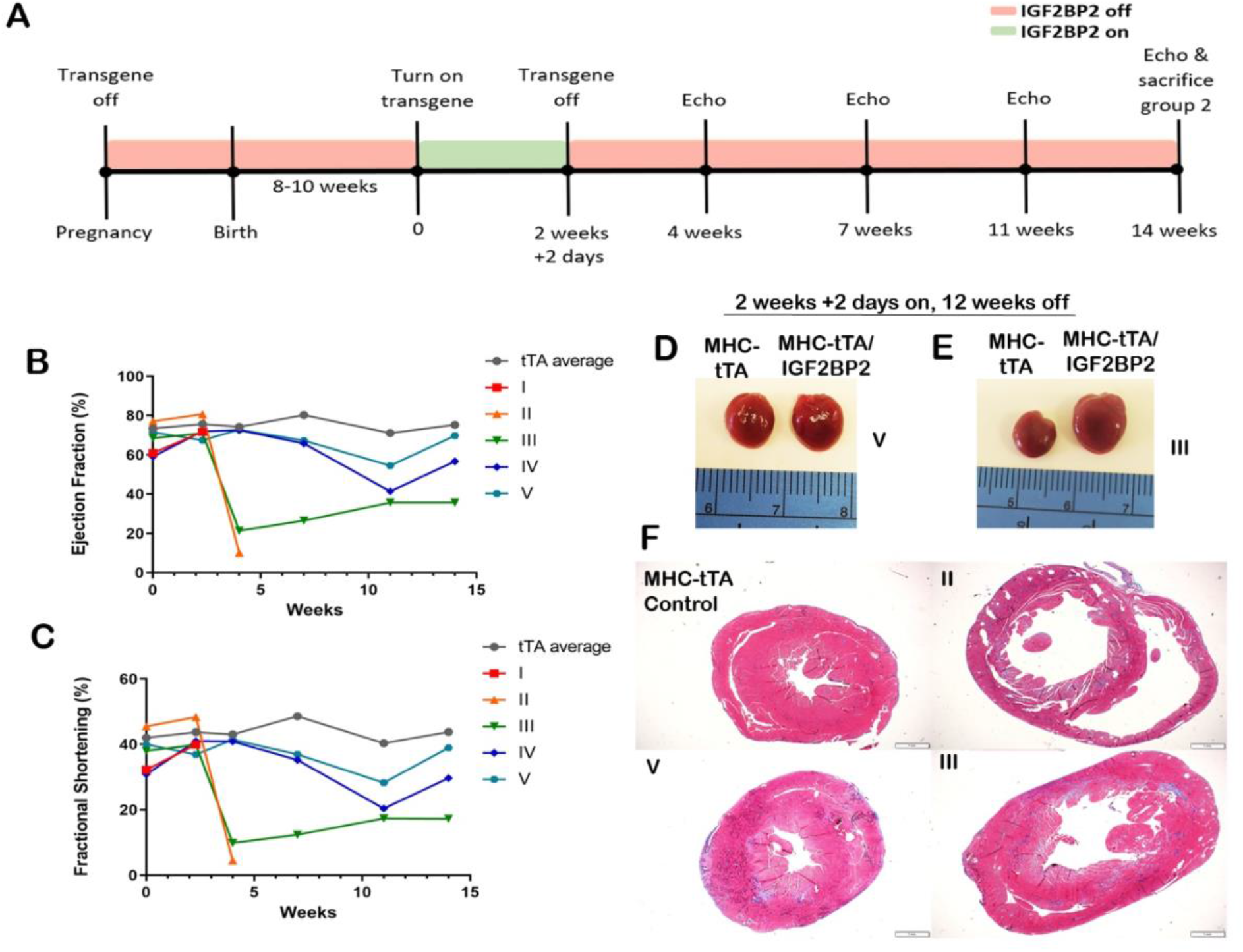
Rescue experiment with only partial recovery. (A) Timeline of experimental design for the experiments described in B-F: transgene on for 2 weeks and 2 days and off for 12 weeks in adult mice. Echocardiograms were performed on all the mice over the course of the experiment and analyzed for Ejection Fraction percentage (B) and Fractional Shortening percentage (C). Three MHC-tTA controls were averaged together and are graphed in gray. The 5 MHC-tTA/IGF2BP2 mice are graphed separately and numbered I-V. (D,E) Picture of hearts at the end of the 2 weeks and 2 days transgene induction followed by 12 weeks of being off: (D) MHC-tTA control heart (left) and an MHC-tTA/IGF2BP2 heart (right) with normal EF% (mouse no.V) and (E) MHC-tTA control heart (left) and an MHC-tTA/IGF2BP2 heart (right) with impaired EF% (mouse no.III). (F)Trichrome staining of an MHC-tTA control and the different outcomes of the MHC-tTA/IGF2BP2 mice at the end of the experiment. Scale bar is 1mm.

### Major protein changes are found in sarcomeres and mitochondria

To elucidate the underlying mechanisms through which IGF2BP2 helps mediate dilated cardiomyopathies, we prepared protein extracts from hearts expressing either the IGF2BP2 transgene or the Tet transactivator alone for 15 days and analyzed them by mass spectrometry (MS; Fig. S3A, B; Dataset S1). Unsupervised hierarchical clustering identified two distinct groups corresponding to the tTA control and IGF2BP2 transgene-expressing mice (Fig. S3C; Dataset S1). Supervised hierarchical clustering of differentially expressed genes showed a significant upregulation of 587 proteins and significant downregulation of 440 proteins (FDR=0.05, S=0.1) (Fig.5A). Among the proteins that were significantly affected by hIGF2BP2 expression, 71 of them (30 downregulated and 41 upregulated) are associated with contractile proteins, as defined in Gene Ontology – Cell Components (GO:0043292).

At least 115 genes have been associated with various familial DCM syndromes (Lehnart & Wehrens, 2022; Maggi *et al*, 2021; McNally & Mestroni, 2017; Rosenbaum *et al*, 2020; Stege *et al*, 2021). Of these, 28 genes are associated with sarcomeres in general or Z discs in particular. Strikingly, 18 of these sarcomeric genes encode proteins significantly downregulated by hIGF2BP2 expression in the transgenic mice, and 10 of these are considered definitive for the DCM phenotype (Table 1). These results correlate nicely with the pathology and physiology of the transgenic hearts, further supporting the evidence that expression of IGF2BP2 in the heart causes DCM.

**Table 1.** Genes associated with DCM and their regulation by IGF2BP2 expression

| Gene code | Gene name | Cellular location | Regulation by IGF2BP2 | Reference |
| --- | --- | --- | --- | --- |
| **LMNA** | Lamin A/C | Nuclear envelope |  | Pereira, 2020 |
| **RBM20** | RNA-binding protein 20 | Nuclear |  | Pereira, 2020 |
| **TTN** | Titin | Sarcomere | down | Pereira, 2020 |
| **MYH7** | Myosin-7 | Sarcomere |  | Pereira, 2020 |
| **MYH6** | Myosin-6 | Sarcomere | down | Pereira, 2020 |
| **TNNT2** | Troponin T, cardiac muscle | Sarcomere | down | Pereira, 2020 |
| **MYBPC3** | Myosin-binding protein C, cardiac-type | Sarcomere | down | Pereira, 2020 |
| **TNNI3** | Troponin I, cardiac muscle | Sarcomere | down | Pereira, 2020 |
| **TPM1** | Tropomyosin alpha-1 chain | Sarcomere | down | Pereira, 2020 |
| **MYL2** | Myosin regulatory light chain 2 | Sarcomere | down | Pereira, 2020 |
| **MYL3** | Myosin light chain 3 | Sarcomere | down | Pereira, 2020 |
| **ACTC1** | Alpha-cardiac actin | Sarcomere |  | Pereira, 2020 |
| **ACTN2** | Alpha-actinin 2 | Z-disc |  | Pereira, 2020 |
| **ANKRD1** | Ankyrin repeat domain-containing protein 1 | Z-disc |  | Pereira, 2020 |
| **CSRP3** | Cysteine and glycine-rich protein 3 | Z-disc | down | Pereira, 2020 |
| **LDB3** | LIM domain-binding protein 3 | Z-disc | down | Pereira, 2020 |
| **MYOZ2** | Myozenin-2 | Z-disc | down | Pereira, 2020 |
| **TCAP** | Telethonin | Z-disc | down | Pereira, 2020 |
| **VCL** | Vinculin | Z-disc | up | Pereira, 2020 |
| **MYPN** | myopalladin | Z-disc |  | Pereira, 2020 |
| **DES** | Desmin | Intermediate filament |  | Pereira, 2020 |
| **FLNC** | Filamin-C | Intermediate filament | up | Pereira, 2020 |
| **SYNM** | Synemin | Intermediate filament |  | Maggi, 2021 |
| **BAG3** | Co-chaperone 3 | Intracellular chaperone |  | Pereira, 2020 |
| **DSP** | Desmoplakin | Intercalated disc | up | Pereira, 2020 |
| **PKP2** | Plakophilin 2 | Intercalated disc |  | Pereira, 2020 |
| **TAZ** | Tafazzin | Mitochondrial membrane |  | Pereira, 2020 |
| **DTNA** | Dystrobrevin-alpha | Plasma membrane |  | Pereira, 2020 |
| **SCN5A** | Alpha-subunit of the cardiac sodium channel | Plasma membrane |  | Pereira, 2020 |
| **JPH2** | Junctophilin-2 | Sarcoplasmic reticulum, plasm membrane | down | Wehrens,2022 |
| **PLN** | Phospholamban | Sarcoplasmic reticulum |  | Sillje, 2021 |
| **CRYAB** | Alpha-crystallin B chain | Nucleus, cytoplasm, membrane | down | Sillje, 2021 |

A GO analysis of the biological processes and cellular components of the proteins significantly downregulated by IGF2BP2 expression (Fig. 5B and Dataset S1) reveals a strong correlation with mitochondrial (309/440=70%, FDR=2.04×10^−216^) and metabolic processes (315/440=72%, FDR=4.54×10^−55^). The same analysis of the upregulated proteins also shows a strong correlation with metabolic processes, albeit different proteins (Fig. 5C and Dataset S1). However, while a similar number and percentage of the upregulated proteins are associated with metabolic processes (359/587=61%, FDR=2.82×10^−41^), only 98 of the upregulated proteins are associated with mitochondria (17%). To gain insight into why some of the proteins associated with metabolic processes are downregulated while others are upregulated by IGF2BP2 expression, we correlated mitochondrial association with metabolic processes for each group. As seen in Figure S4, over 75% of the downregulated metabolic processes proteins are mitochondrial, as opposed to only 20% of the upregulated proteins. This highly significant correlation (χ^2^<0.0001) indicates that IGF2BP2 expression in the heart causes mitochondrial proteins, and, as a result, mitochondrial metabolism, to be greatly diminished; non-mitochondrial metabolic pathways, in contrast, are upregulated in an attempt to compensate for the heart’s metabolic requirements. Several up-and down-regulated proteins were validated by western blot analysis (Fig.5D, E).

**Figure 5.**
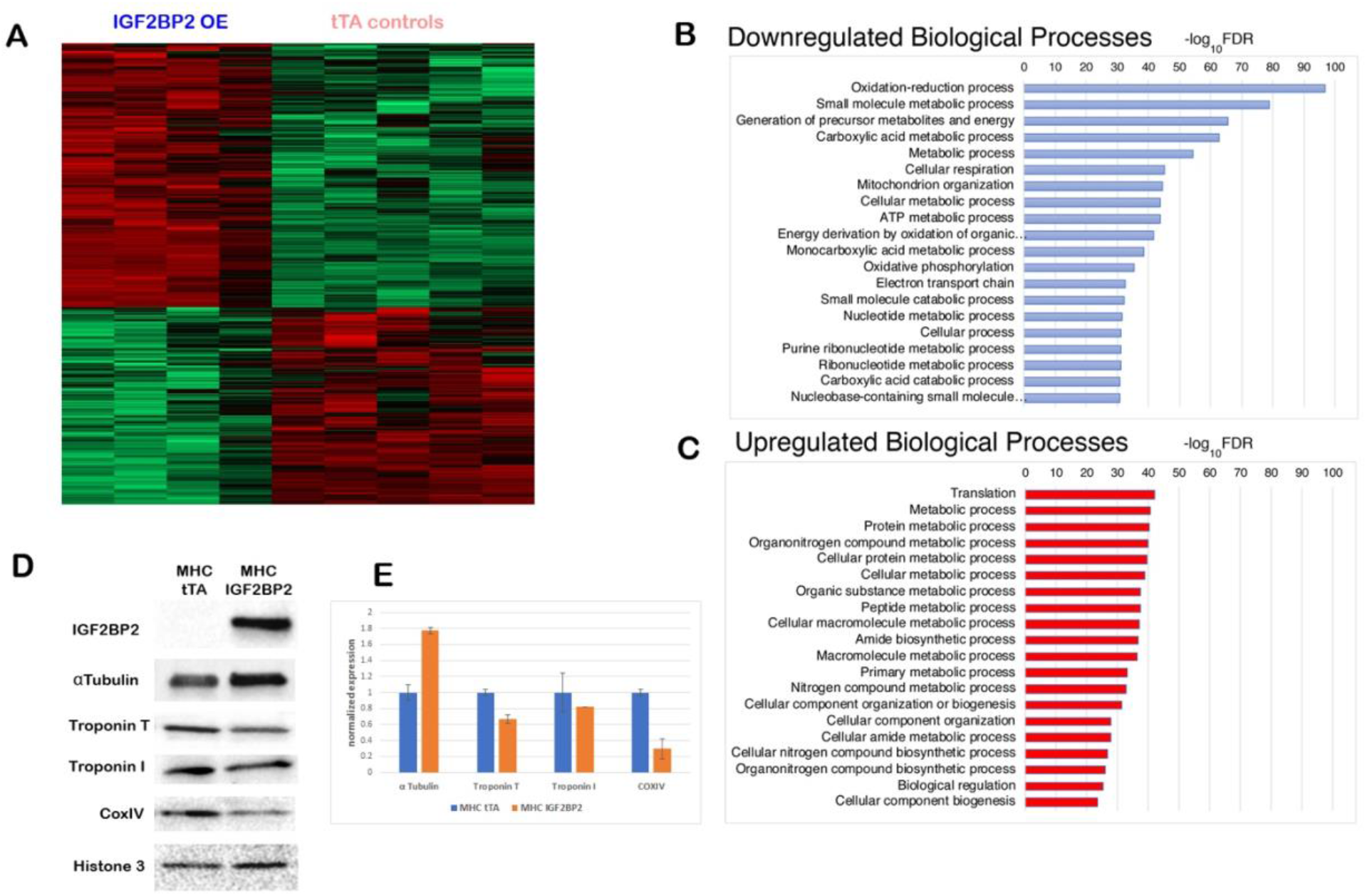
Proteomics of uninduced and induced adult hearts. Protein extracts were prepared from MHC-tTA adult mice hearts (5 samples, tTA controls) or from same-age siblings MHC-tTA/IGF2BP2 mice in which IGF2BP2 expression was activated for 15 days (4 samples, IGF2BP1 OE) and analyzed by mass spectrometry. (A) Supervised hierarchical clustering of all 9 samples of differentially expressed proteins (with Z score, FDR=0.05, S=0.1; red, downregulated; green, upregulated). Top 20 Biological processes in GO analysis were ranked by -log_10_FDR (false discovery rate) of downregulated (B) and upregulated (C) proteins. (D) Western blot validation of proteins shown to be up-(IGF2BP2) or down-regulated (Tubulin, Troponin T, Troponin I, and COXIV) by MS analysis mass spectrometry. Histone 3 was used as a loading control. Quantification of the western blot analysis with two biological samples in each group ±SEM.

To analyze the effect of IGF2BP2 on sarcomere structure at the cellular level, IGF2BP2 expression was activated in primary cardiomyocytes for 7 days. Cells were stained with anti-IGF2BP2, anti-MF20 (a pan myosin heavy chain antibody), and DAPI (Fig. S5A), and images were analyzed using an algorithm in ImageJ which highlights the direction of polarized pixels, based on coherency mapping (Puspoki *et al*, 2016). As seen in Fig. S5B, while control cardiomyocytes reveal a bright MF20 staining upon coherency mapping, indicating a more global orientation of myofibrils within the cell, IGF2BP2-expressing cardiomyocytes lose organization of pixels in the MF20 staining, and exhibit a duller, more spotted appearance.

These results suggested that changes in cardiomyocyte structure upon IGF2BP2 induction might also be apparent at the ultrastructural level. Transmission electron microscopy (EM) images were acquired from the LV of adult hearts expressing either MHC-tTA alone or MHC-tTA/IGF2BP2, 2 weeks after activating transgene expression (Fig 6A). In accordance with the proteomics and myofibril staining data, the sarcomeres of the tTA-IGF2BP2 heart showed an altered structure, with the distance between the Z lines elongated (sarcomere length) and the width of the myofibrils shortened (Fig 6B-C), a phenotype observed in dilated cardiomyopathy (DCM) (Collins *et al*, 2021; Wu *et al*, 2021). Western blot analysis shows an approximately 30% reduction in the sarcomere Z disk-localized αActinin protein in hIGF2BP2 expressing mice (Fig. S6A, B). Interestingly, the RNA for both αActinin2 and several mitochondrial proteins are actually upregulated in primary cardiomyocytes after 5 days of hIGF2BP2 expression (Fig. S6C), suggesting that IGF2BP2 may be regulating sarcomeric and mitochondrial genes by binding their RNA transcripts, stabilizing the mRNA, but inhibiting translation. A similar repressive role in the translation of CDC42 mRNA in the heart has recently been attributed to enhanced IGF2BP2 binding (Qian *et al*, 2021).

**Figure 6.**
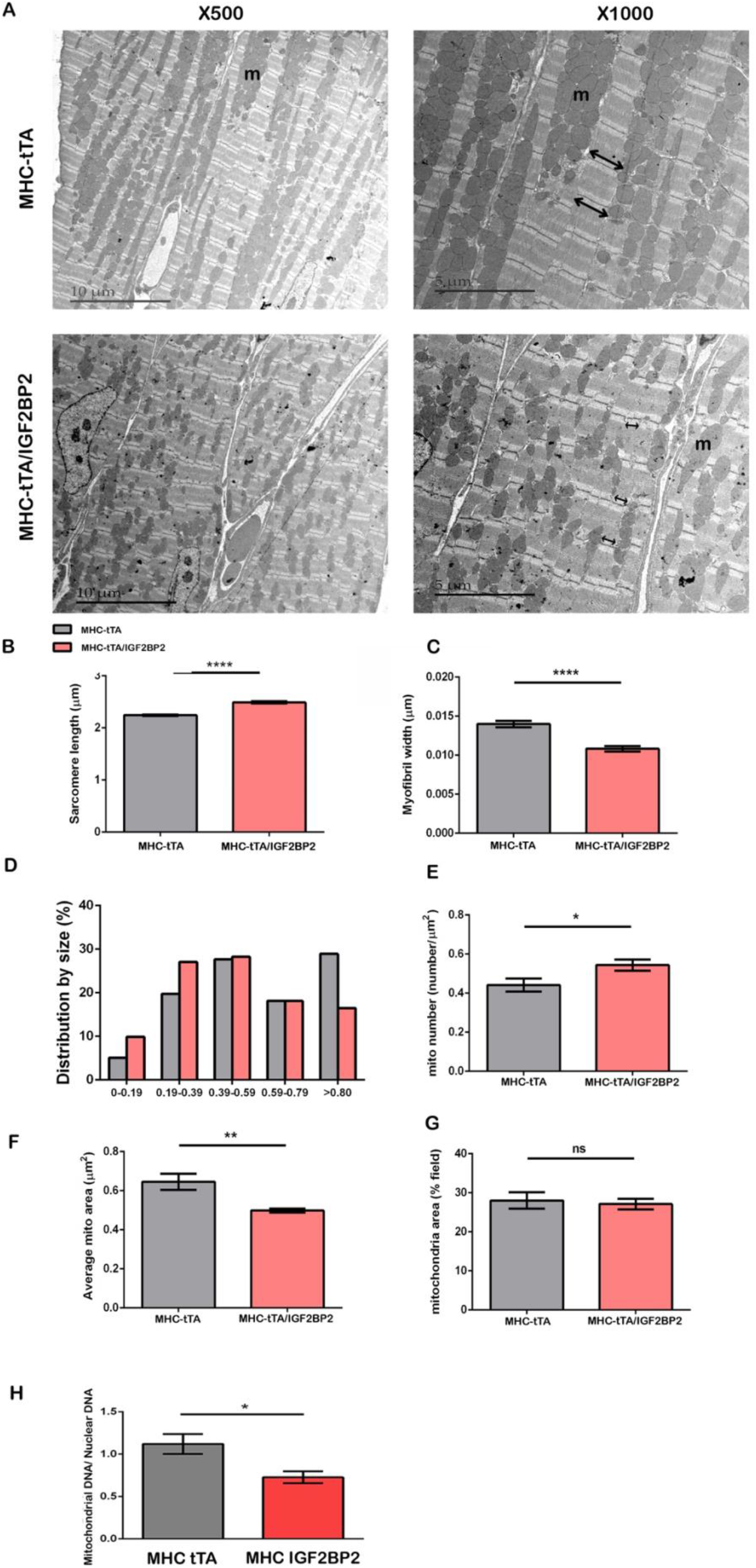
EM images of the myocardium from transgenic mice. (A) Representative EM images of the LV myocardium of mice. Left panels, magnification was x500, and right panels, x1000. “m” indicates mitochondria. Note the absence of the H zone in both the MHC-tTA and MHC-tTA/IGF2BP2 hearts, indicating that both are in contraction. The double headed arrows indicate the width of the myofibrils in the myocardium. Activation of IGF2BP2 expression leads to increased sarcomere length (B; *p* < 0.001) and decreased myofibril width (C; *p* < 0.001). IGF2BP2 activation also causes fractionalization of mitochondria, leading to an accumulation of smaller mitochondria (D) with a concomitant increase in their number per field (E; *p*=0.0358). Accordingly, the average size per mitochondrion drops (F; *p*=0.0037), although mitochondria still make up the same percentage of the area of the total field (G). (H) Mitochondrial DNA is reduced in hearts expressing IGF2BP2, as measured by PCR (*p*=0.0466). \**p* <0.05, \*\**p*<0.01 and \*\*\*\**p* <0.001.

The proteomics data predicted that IGF2BP2 expression should have a strong effect on the mitochondria in transgenic hearts. Indeed, upon IGF2BP2 overexpression, a disordered alignment of mitochondria is observed in EM grids, along with alterations in shape and size (Fig 6A, D). Quantitative analysis shows that the number of mitochondria is 19% higher in the IGF2BP2-induced hearts (Fig 6E), although the average size was reduced by 23% (Fig 6F). Given that the total mitochondrial area per field is the same in both the control and overexpression hearts (Fig. 6G), these results indicate that the mitochondria are fragmented and not undergoing fusion properly when IGF2BP2 expression is activated. In a rat coronary artery constriction model, heart failure was shown to cause a similar phenotype of increased numbers of fragmented, smaller mitochondria (Chen *et al*, 2009). OPA1 is a mitochondrial inner membrane protein that helps regulate fusion of mitochondria and is downregulated by ischemia and in heart failure. Knockdown of OPA1 causes fragmentation of mitochondria in cultured cardiomyocytes (Chen *et al*., 2009), and, in mice, leads to heart failure (Wai *et al*, 2015). Notably, OPA1 is one of the mitochondrial proteins downregulated after IGF2BP2 activation (Dataset1). It is interesting to note that OPA1 is also an essential protein for initiating mtDNA replication and maintaining mtDNA integrity (Elachouri *et al*, 2011). The abundance of a mitochondrial DNA sequence was compared to that of a nuclear genomic DNA sequence in DNA extracted from the same hearts used for the MS analysis. We found that there was almost 40% less mtDNA in the IGF2BP2-induced hearts than in the MHC-tTA controls (Fig. 6H). Taken together, these results confirm that overexpression of IGF2BP2 in the heart causes a profound impairment of both sarcomere and mitochondrial morphology.

### IGF2BP2 expression in patients after MI or with DCM

Having observed that hIGF2BP2 expression in mouse hearts is sufficient to cause dilated cardiomyopathy and that IGF2BP2 RNA is up-regulated in hearts that have undergone cardiac stress, we wondered whether IGF2BP2 protein expression is elevated in human pathologies. Heart tissue from patients with various cardiac diseases was stained for IGF2BP2 protein (Fig. 7). While IGF2BP2 expression was not observed in hypertrophic heart samples, IGF2BP2 expression was indeed detected in both myocardial infarction (MI; n=13) and DCM (n=20) hearts, with expression significantly higher in DCM as compared to MI sections (p=0.0404). These findings in human patients correlate well with the results from the mouse experiments, showing that IGF2BP2 is up-regulated upon cardiac stress and helps to remodel the heart.

**Figure 7.**
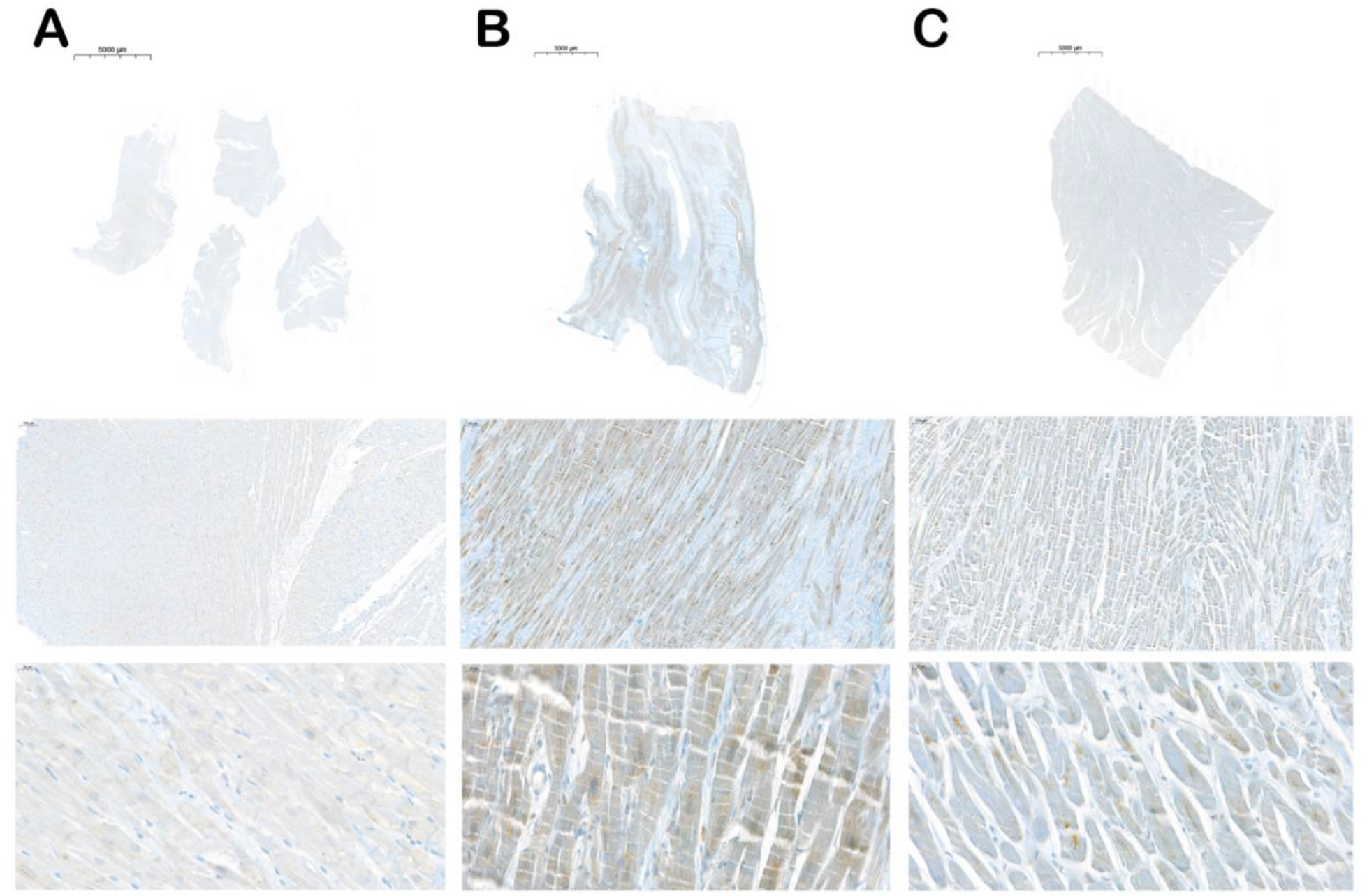
Expression of IGF2BP2 in human cardiomyopathies. Human pathological heart sections were stained with antibody against IGF2BP2. (A) Hypertrophic phenotype. (B) Dilatative cardiomyopathy. (C) Myocardial infarction. First row shows overview images at 0.3x magnification, scale bar 5000µm, the second row shows images at 8x magnification, scale bar 100µm, and the third row shows detail images at 40x magnification, scale bar 20µm.

## Discussion

Here we demonstrate for the first time the involvement of IGF2BP2 in response to cardiac stress and in the development of cardiomyopathies. IGF2BP2 is normally expressed in the heart during embryogenesis, and its expression is maintained at a low level postnatally into adult life. Expression of transgenic IGF2BP2 in the mouse heart, either from birth or in the adult, causes an enlarged, remodelled, malfunctioning heart that leads to death within a period of approximately 3 to 4 weeks. Strikingly, IGF2BP2 is upregulated not only in mouse models of cardiac ischemia and pressure overload, but also in human patients with dilated cardiomyopathy and after myocardial infarction (Fig. 7). In several mouse models of induced cardiac stress, rescue of the heart and recovery of normal morphology is correlated with downregulation of IGF2BP2, raising the possibility that upregulation of IGF2BP2 under stress conditions may initially be a compensatory response that later needs to be restricted to enable full recovery.

This hypothesis would predict that turning off IGF2BP2 expression in a timely fashion could reverse the DCM phenotype it induced. Using a conditional mouse model, IGF2BP2 was upregulated in adult hearts for 2 weeks and then turned off. Although increased expression of IGF2BP2 for 2 weeks causes a DCM phenotype with the associated morphological, structural changes in the myocardium and in gene expression, these changes were not severe enough to impair heart function. In addition, all these mice ultimately recovered, and their hearts returned to normal size. Increasing the overexpression period by only two days (16 as opposed to 14 days) caused a significant change in the phenotype of the mice: two mice died within two weeks of turning off hIGF2BP2, one mouse survived until the end of the experiment (12 weeks) but had an enlarged, malfunctioning heart, and only two mice were healthy three months post-abrogation of hIGF2BP2. These results indicate that, in mice, IGF2BP2 upregulation triggers a spiral into cardiac disease that is initially reversible but becomes independent of IGF2BP2 expression after 2 weeks.

The mechanisms by which overexpression of IGF2BP2 leads to DCM and death appear to be multiple and complex. In our proteomics analysis, over 300 mitochondrial proteins were significantly reduced by hIGF2BP2 expression, representing by far the highest ranking Cellular Component GO group influenced by IGF2BP2. The short, fragmented mitochondria phenotype observed in the EM sections of the hIGF2BP2 expressing hearts is fully consistent with the broad downregulation of mitochondrial proteins and appears very similar to the mitochondria seen in ischemic heart failure (Chen *et al*., 2009). Mitochondria constitute approximately 40% of the cytoplasm of an adult cardiomyocyte (Li *et al*, 2020), and mitochondrial dysfunction has long been known to be associated with the development of heart disease (Zhou & Tian, 2018). Because many of the downregulated proteins are also classified as metabolic, such a largescale assault on mitochondrial proteins can explain why these hearts proceed to heart failure. Accordingly, energy must be generated in these mice via alternative pathways, and this compensation is indeed observed as an upregulation of many other non-mitochondrial, metabolic proteins. A very similar phenotype, involving downregulation of aerobic mitochondrial metabolism genes and an upregulation of glycolytic glucose metabolism genes, is observed in mice induced to develop DCM by a mutation in phospholamban (Burke *et al*, 2016a). Similarly, genetic mitochondrial diseases, where the function of the mitochondria is compromised by mutations in mitochondrial proteins, are associated with cardiac abnormalities as well. Electrocardiographic and/or echocardiographic abnormalities are found in about 30% of adults with mitochondrial disease (Duran *et al*, 2019; El-Hattab & Scaglia, 2016; Orsucci *et al*, 2021; Pankuweit & Richter, 2015). Indeed, improving mitochondrial function is considered an emerging approach for treating heart failure (Brown *et al*, 2017).

Recently, it was shown that IGF2BP2 associates with αActinin2 protein in murine primary cardiomyocytes, where it helps mediate intracellular localization of RNAs encoding mitochondrial proteins to the vicinity of the sarcomere (Ladha *et al*., 2021). We also observed association of IGF2BP2 with mitochondrial RNAs, leading to an increased level of these transcripts. Nevertheless, mitochondrial proteins in the hIGF2BP2 expressing hearts are downregulated, indicating that IGF2BP2 is likely repressing translation of many of these bound RNAs. IGF2BP2 has been shown to repress translation in several different settings, including translation of mitochondrial UCP RNA in brown fat (Dai *et al*, 2015). Significantly, these authors observed that over 40% of the RNAs associated with IGF2BP2 (as seen by immunoprecipitation) are mitochondrial. OPA1 mRNA, which encodes a mitochondrial inner membrane fusion protein that causes mitochondrial fragmentation when downregulated, is one such target of IGF2BP2 (Conway *et al*., 2016), and its protein product is also downregulated in the IGF2BP2-overexpressing hearts (Dataset S1). Thus, it appears that IGF2BP2 overexpression represses the expression of a large number of mitochondrial RNAs, likely inhibiting both OPA1-mediated fusion and mitochondrial function.

The proteomic data also show that many sarcomeric proteins are reduced by elevated IGF2BP2 expression. Here, as with mitochondrial RNAs, IGF2BP2 appears to bind RNAs encoding these proteins, protecting the RNAs from degradation but repressing their translation. Many of the downregulated sarcomeric or Z-disc associated proteins were previously identified as genes associated with DCM in patients. Consistent with these findings, primary cardiomyocytes overexpressing IGF2BP2 show a loss of directionality and polarization. At the EM level, the sarcomeres of hearts overexpressing IGF2BP2 are elongated and thinner, a phenotype observed in DCM hearts. Taken together, these data suggest that IGF2BP2 upregulation impairs the translation of many IGF2BP2 target RNAs, particularly in mitochondria and sarcomeres, leading to DCM. Future studies should help clarify which mechanisms are responsible for the observed phenotypes.

In recent years, RNA methylation has been observed to play a role in cardiomyophathies. m^6^A RNA methylation is upregulated when heart failure occurs in both humans and mice, as well as in primary cultured cardiomyocytes exposed to ischemia (Mathiyalagan *et al*, 2019). When RNA methylation is upregulated either by overexpression of the methylation writer, METTL3A, or downregulation of the methylation eraser, FTO, hearts become hypertrophic (Dorn *et al*, 2019) and show impaired cardiac contractility (Mathiyalagan *et al*., 2019). Conversely, mice in which METTL3A is knocked down do not become hypertrophic upon aging or when exposed to cardiac stress (Dorn *et al*., 2019), and MI-induced cardiac injury can be rescued by overexpression of FTO (Mathiyalagan *et al*., 2019). Recently, the IGF2BP proteins have been proposed to be readers of m^6^A RNA methylation, i.e., methylation of their RNA targets enhances binding by the proteins (Huang *et al*., 2018). It is tempting to speculate that increased m^6^A RNA methylation could lead to elevated RNA binding by IGF2BP2, facilitating its role in mediating cardiomyopathies. In this regard, it is interesting to note that METTL3 RNA has been reported to be itself a target of IGF2BP2 in HEK293 cells (Huang *et al*., 2018). In addition, IGF2BP2 has recently been shown to cooperate with miR-133a and AGO2 in an RNA methylation-dependent manner; increased m^6^A methylation leads to increased IGF2BP2 binding of miR-133a and recruitment of AGO2 to target RNAs, causing translational repression and cardiac hypertrophy (Qian *et al*., 2021). Future experiments will more fully determine how IGF2BP2 may interact with the methylation machinery in the heart.

In summary, we propose a model in which cardiac stress upregulates IGF2BP2, which in turn induces cardiac remodelling that acts as an adaptive response, allowing the heart to continue to function. This activation causes a shift in the heart from mitochondrial-to non-mitochondrial based metabolism, as well as a disruption of normal sarcomeric organization. The resulting dilated cardiomyopathy eventually leads to heart fatigue and death, unless IGF2BP2 expression is reduced in time. We propose that IGF2BP2 may be a useful target for the treatment of cardiac disorders in which there are upregulated levels of the protein.

## Materials and Methods

### Mice

#### Transgenic animals

All animal procedures were performed in accordance with the local animal ethics committee (MD-17-15389, MD-20-16134).

The generation of LSL-GFP-IGF2BP1 (VICKZ1) mice has been previously described (Rosenfeld *et al*, 2019) These mice were crossed with either PKC-CRE mice that were available in our facility or with αMHC-CRE mice, a kind gift from Prof. Eldad Tzahor, Weizmann Institute.

TRE-human p62 (IGF2BP2) mice were a kind gift from Alexandra K. Kiemer, Saarland University, and have been previously described (Tybl *et al*, 2011). In short, the human IGF2BP2 protein (p62) transgene is under control of the trans responsive element cytomegalovirus (TRE-CMVmin) promotor. In order to induce human IGF2BP2 expression in the heart, transgenic mice were bred with αMHC-tTA mice, a kind gift from Prof. Eldad Tzahor, Weizmann Institute, which carry the gene for tetracycline transactivator (tTA) under a cardiac specific promoter (αMHC). Cardiac-specific expression of the transgene can be switched off by the application of tetracycline or switched on by the withdrawal of the tetracycline. To repress human IGF2BP2 expression during embryonic and early postnatal heart development, mothers were given 0.5mg/ml tetracycline (Tamar Inc., Israel) with 3% sucrose in the drinking water. Mice were then kept on tetracycline until induction at 8-10 weeks of age. All heart weight to body weight (HW/BW) or tibia length to body weight (TL/BW) ratios were done per sex. In PGK IGF2BP1, MHC IGF2BP1, Rescue IGF2BP2 are all heart weights of male mice. MHC IGF2BP2 in the adult data and mass spectrometry are taken from female mice. All echocardiograms, the rescue experiment and the electron microscopy were done on male mice.

### Transverse Aortic Constriction (TAC)

TAC mice models (n=3) to induce chronic pressure overload of the left ventricle (deAlmeida *et al*., 2010; Rockman *et al*, 1991) were performed by standard protocol in the Kehat lab at the Technion Institute (Israel).

### Echocardiography

Short axis measurements were obtained for heart function evaluation, by transthoracic echocardio-graphy performed on isoflurane-sedated mice using a Vevo LAZR-X VisualSonics device for IGF2BP2 mice and the VEVO700 for the IGF2BP1 mice. Three frames of short axis were analyzed for three Diastole and systole cycles. All echocardiography measurements were performed in a blinded manner. Data is presented as mean + SEM. ** p<0.01,****p<0.0001; statistical significance was calculated using one-way ANOVA followed by Dunnett’s post hoc test relative to tTA group.

### Histology staining and IHC

Hearts were fixed and paraffin embedded. Sections of 5µm thickness were then processed for tissue staining. H&E and Trichrome staining were done using standard protocol. Collagen staining to identify fibrosis was done using standard Sirius Red protocol. Immunohistochemistry was performed using the primary antibodies summarized below:

| Primary ab | Source |  |  |  |  |
| --- | --- | --- | --- | --- | --- |
| Protein | Rabbit | Dilution western | Dilution ICF | Company | Cat no. |
| IGF2BP2 | Rabbit | 1:3000 | 1:250 | Sonja Kessler |  |
| $\alpha/\beta$ Tubulin | Mouse | 1:3000 | | Cell Signaling | 2148s |
| $\alpha$ Tubulin | Mouse | 1:1000 | | Cell Signaling | 3873 |
| Troponin T | Mouse | 1:80 |  | Hybridoma bank |  |
| Troponin I | Mouse | 1:80 |  | Hybridoma bank |  |
| MF20 | Mouse | 1:400 | 1:150 | Hybridoma bank |  |
| COXIV | Mouse | 1:10,000 |  | abcam | ab33985 |
| Histone 3 | Rabbit | 1:1000 |  | Cell Signaling | 9701 |
| $\alpha$ Actinin | | | | Cell Signaling | 6487s |
| Secondary ab |  |  |  |  |  |
| DAPI | Goat anti Mouse |  | 1:1000 | Biorad |  |
| 488 | Donkey anti Rabbit |  | 1:400 | Jackson immuno research |  |
| Cy3 |  |  | 1:400 | Jackson immuno research |  |

Autofluorescence in immunofluorescence staining was quenched using the Vector® TrueVIEW® Autofluorescence Quenching Kit (Vector Laboratories, SP-8400-15) following manufacturer’s instructions. Wheat germ-agglutinin staining (1 mg/ml, Sigma) was performed for 1 hour, treated with Vector TrueVIEW kit, stained with DAPI and mounted with VECTASHIELD Vibrance Antifade Mounting Medium (Dorn *et al*., 2019). In enzymatic reactions, the ImmPRESS® HRP Horse Anti-Rabbit IgG Polymer Detection Kit, Peroxidase (Vector Laboratories,MP-7401) and the ImmPACT® DAB Substrate, Peroxidase (HRP) (Vector Laboratories, SK-4105) were used.

Hearts were cut in half in adult, taking the apex for molecular analysis and a region from under the midline to base for sectioning. Sections prepared for experiments of IGF2BP2 expression from birth were prepared from the apex to base region. 3D modelling was done using High-Resolution Episcopic Microscopy (HREM), as previously published (Pokhrel *et al*, 2022).

### Electron Microscopy

For perfusion, mice were weighed and anesthetized by i.p. injections with ketamine/xylazine solution (50 mg/kg ketamine/7.5 mg/kg xylazine in 0.9% NaCl solution). The animals were transcardially pre-perfused with ice cold 10ml of 0.1 M phosphate buffer, followed by perfusion with 10ml ice cold fresh prepared Karnovsky fixative containing 2% paraformaldehyde and 2.5 % glutaraldehyde EM grade, in 0.1M sodium cacodylate buffer pH 7.3.

Tissue was washed 4 times with sodium cacodylate and postfixed for 1 hr with 1% osmium tetroxide and 1.5% potassium ferricyanide in sodium cacodylate, and washed 4 times with the same buffer, followed by dehydration with a graded ethanol series, followed by 2 changes of propylene oxide. Cells were then infiltrated with a series of epoxy resin and polymerized in the oven at 60°C for 48 hours. 80 nm sections were obtained and stained with uranyl acetate and lead citrate. Sections were observed by Jeol JEM 1400 Plus Transmission Electron Microscope, and pictures were taken using a Gatan Orius CCD camera.

### Human sections

Immunohistochemical staining was performed with the Ventana Benchmark Ultra System, using the Ultra View Universal DAB Detection Kit (#5269806001, Roche Diagnostics, Rotkreuz, Switzerland) for antibody detection, according to the manufacturer’s instructions. Anti-IGF2BP2/p62 antibody (References : https://doi.org/10.1152/ajpgi.00005.2012 and doi: 10.1016/S0002-9440(10)61770-1) which detects both isoforms was incubated in a 1:100 dilution for 32 min at room temperature. Epitope retrieval was achieved with Cell Conditioning Solution 1 (#5279801001, Roche Diagnostics, Rotkreuz, Switzerland) at 95°C for 64 min. Samples were examined by an independent investigator blinded to experimental conditions. The intensity of each section was scored (intensity 0 = no detectable staining, 1 = weak staining, 2 = moderate staining, and 3 = strong staining). Statistics were done using Chi2 test.

Samples used in this project were with the permission of the local ethics commission (EK number 4077).

### Cardiomyocyte isolation and Immunofluorescence staining

Cardiomyocytes were isolated using Neonatal Cardiomyocyte Isolation System (#LK003300; Worthington Biochemical Corporation) following manufacturer’s instructions. Cardiomyocytes were seeded with DMEM-F12 (Biological Industries, Israel) with 10% FCS and 1% ciprofloxacin (Bayer) for 48 hours and then grown in DMEM-F12 with 0.1% serum and 1% ciprofloxacin for 3-4 days on cover glasses coated with BME (Celtrux). Cells were fixed with 4% formaldehyde and incubated with primary and secondary antibodies consecutively (see *Histology staining and IHC* for full list of antibodies).

Cells were imaged using the Nikon Spinning Disk microscope. For directionality of pixels in MF20 cardiomyocyte staining, ImageJ software was used, using the OrientationJ plugin as instructed (Puspoki *et al*., 2016). 3 repeats of the experiment were performed.

### Sample preparation for MS analysis

Heart tissue was homogenized in RIPA buffer containing protease and phosphatase inhibitors, clarified by centrifugation, and the supernatant was subjected to protein precipitation by the chloroform/ methanol method (Wessel & Flugge, 1984). The precipitated proteins were solubilized in 100 μl of 8M urea, 10 mM DTT, 25 mM Tris-HCl pH 8.0 and incubated for 30 min at 22°C. Iodoacetamide (55 mM) was added followed by incubation for 30 min (22°C, in the dark), followed by re-addition of DTT (10 mM). 25 μg of protein was transferred into a new tube, diluted by the addition of 7 volumes of 25 mM Tris-HCl pH 8.0 and sequencing-grade modified Trypsin (Promega Corp., Madison, WI) was added (0.4 μg/ sample) followed by incubation overnight at 37°C with agitation. The samples were acidified by addition of 0.2% formic acid and desalted on C18 home-made Stage tips. Peptide concentration was determined by Absorbance at 280 nm and 0.75 µg of peptides were injected into the mass spectrometer.

### nanoLC-MS/MS analysis

MS analysis was performed using a Q Exactive-HF mass spectrometer (Thermo Fisher Scientific, Waltham, MA USA) coupled on-line to a nanoflow UHPLC instrument, Ultimate 3000 Dionex (Thermo Fisher Scientific, Waltham, MA USA). Peptides dissolved in 0.1% formic acid were separated without a trap column over a 120 min acetonitrile gradient run at a flow rate of 0.3 μl/min on a reverse phase 25-cm-long C18 column (75 μm ID, 2 μm, 100Å, Thermo PepMapRSLC). The instrument settings were as described (Scheltema *et al*, 2014). Survey scans (300–1,650 m/z, target value 3E6 charges, maximum ion injection time 20 ms) were acquired and followed by higher energy collisional dissociation (HCD) based fragmentation (normalized collision energy 27). A resolution of 60,000 was used for survey scans and up to 15 dynamically chosen most abundant precursor ions, with “peptide preferable” profiles, were fragmented (isolation window 1.8 m/z). The MS/MS scans were acquired at a resolution of 15,000 (target value 1E5 charges, maximum ion injection times 25 ms). Dynamic exclusion was 20 sec. Data were acquired using Xcalibur software (Thermo Scientific). To avoid a carryover and to equilibrate the C18 column, the column was washed with 80% acetonitrile, 0.1% formic acid for 25 min between samples, as per standard protocol.

Raw data were processed using MaxQuant (MQ) version 1.6.5.0 (Cox & Mann, 2008) and the embedded Andromeda search engine (Cox *et al*, 2011). The bioinformatics was performed with Perseus suite (version 1.6.2.3). The data were filtered for reverse, contaminants and identified by site. Then the data were filtered such that a protein had to have non-zero LFQ intensity in all 9 samples with 3 or more peptides. The significantly enriched proteins were found (two-sample *t* test with a permutation-based FDR method) and further selected using an adjusted *p* value <0.05 and S=0.1. Log2-transformed individual values or triplicate means were z-score-normalized prior to hierarchical clustering. Gene Ontology annotation performed using STRING site version 10.5, and the statistical values by the site have been reported. A file containing lists of the proteins at the different stages of filtering has been included as Expanded View data (Dataset S1). The mass spectrometry proteomics data have been deposited in the ProteomeXchange Consortium via the PRIDE (Perez-Riverol *et al*, 2022) partner repository with the dataset identifier PXD037623.

### Gene omnibus (GEO) data sets

Publicly available Gene omnibus (GEO) data sets from murine myocardial infarction samples from mice that underwent ligation of the left coronary artery (GDS488), heart failure due to expression of mutated estrogen receptor-coupled Cre under transcriptional control of cardiac-specific myosin heavy chain (MerCreMer; GDS5469), and induced cardiomyopathy due to exposure to isoproterenol (GDS3596), were analysed for IGF2BP2 expression.

### Statistics

Data are shown as mean ± SEM unless otherwise stated. One-way analysis of variance (ANOVA) was used to determine statistical significance for experiments with more than two groups followed by Dunnett’s post hoc test. Figures with ANOVA analysis where applicable are indicated in corresponding figure legends. Comparison between two groups were carried out using GraphPad software with an unpaired Student’s t-test. In human IHC statistics was performed using a chi-squared test. P-values <0.05 were considered statistically significant and assigned in individual figures.

### Real-time and quantitative polymerase chain reaction

Total RNA from heart tissue was extracted with the NucleoSpin RNA Mini Kit with DNase (Macherey-Nagel). cDNA was synthesized using oligo dT with the Flex Script cDNA Synthesis kit (Quanta bio). qPCR was done on an CFX384 Touch™ Real-Time PCR Detection System (Bio-Rad) using the Fast SYBR Green Master Mix (Applied Biosystems, ThermoFisher Scientific). Results were calculated using Bio-Rad CFX software. The list of primers used are shown below:

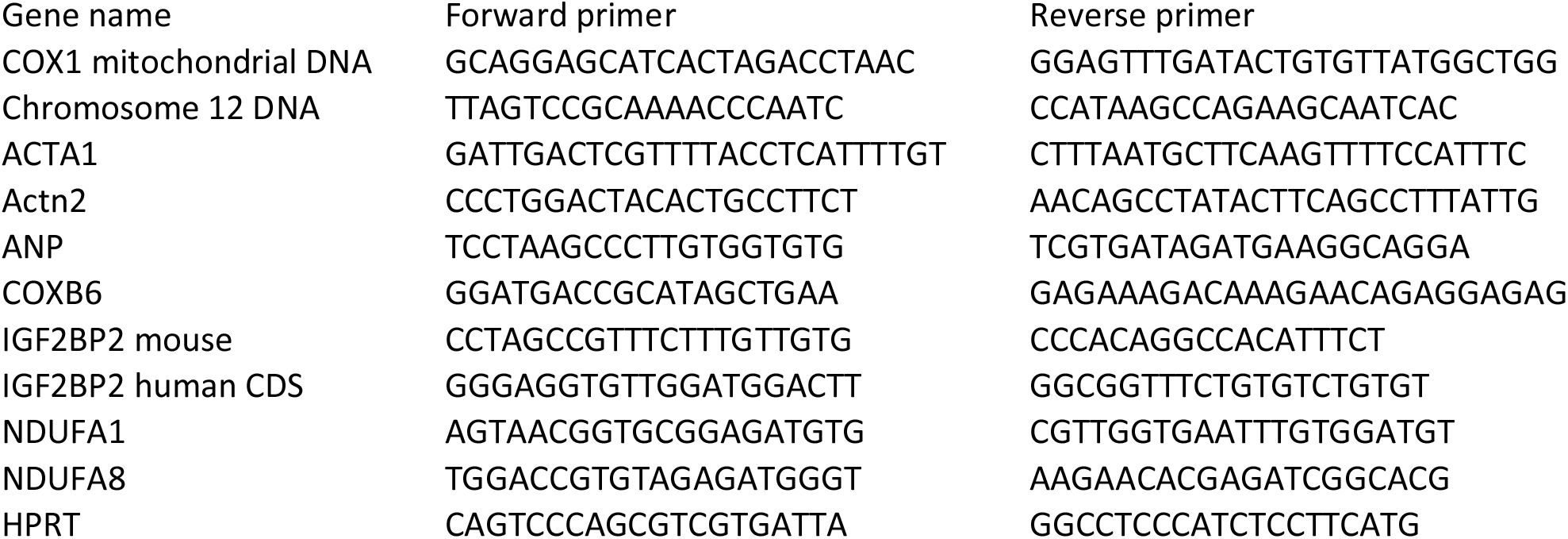

For nuclear and mitochondrial DNA extraction, the Qiagen DNeasy Blood & Tissue Kit (69504) was used. For nuclear DNA, a sequence on chromosome 12 was used as an internal control and a sequence from COX1 was used for mitochondrial DNA.

### Western blot analysis

Tissue extracts were prepared using RIPA buffer with the addition of (β-Glycerophosphate and Sodium orthovanadate) and a protease inhibitor cocktail (Ape Bio). Protein extracts were run on precast SDS-PAGE gels (Bio Rad) and transferred to nitrocellulose membranes prior to imaging with the Bio Rad ChemiDoc Imaging System.

List of antibodies can be found in *Histology staining and IHC*.

## Acknowledgements

Thanks to Dr. Jonathan Balkin for helpful comments, Rinat Abromovitz and all of the Wohl center staff, David Mimon and Tali Benkin, for assistance with the echocardiograms, Nadav Wallis and Hagar Klein for help with mice and cardiomyocyte cultures, Yoav Smith for assistance with the bioinformatics analysis, and Omri from Micha Berger’s lab for the primer sequences for mitochondrial DNA.

This study was funded by the Deutsche Forschungsgemeinschaft (#KE 2519 to J.K.Y. and S.M.K.), the American Friends of the Hebrew University, the Salomon Family Philanthropic Fund of the Hebrew University, and the Saul and Joyce Brandman Cardiovascular Research Hub of the Institute for Medical Research – Israel-Canada (IMRIC) at the Faculty of Medicine of The Hebrew University of Jerusalem.

## Notes

### Competing Interest Statement

The authors have declared no competing interest.

### Summary of Updates

The paper now focuses on the effect of Igf2bp2.

## References

Akerberg AA, Burns CE, Burns CG (2019) Exploring the Activities of RBPMS Proteins in Myocardial Biology. Pediatr Cardiol 40: 1410–1418

Brown DA, Perry JB, Allen ME, Sabbah HN, Stauffer BL, Shaikh SR, Cleland JG, Colucci WS, Butler J, Voors AA et al (2017) Expert consensus document: Mitochondrial function as a therapeutic target in heart failure. Nature reviews Cardiology 14: 238–250

Burke MA, Chang S, Wakimoto H, Gorham JM, Conner DA, Christodoulou DC, Parfenov MG, DePalma SR, Eminaga S, Konno T et al (2016a) Molecular profiling of dilated cardiomyopathy that progresses to heart failure. JCI Insight 1: e86898

Burke MA, Cook SA, Seidman JG, Seidman CE (2016b) Clinical and Mechanistic Insights Into the Genetics of Cardiomyopathy. J Am Coll Cardiol 68: 2871–2886

Carmel MS, Kahane N, Oberman F, Miloslavski R, Sela-Donenfeld D, Kalcheim C, Yisraeli JK (2015) A Novel Role for VICKZ Proteins in Maintaining Epithelial Integrity during Embryogenesis. PloS one 10: e0136408

Chen L, Gong Q, Stice JP, Knowlton AA (2009) Mitochondrial OPA1, apoptosis, and heart failure. Cardiovasc Res 84: 91–99

Collins HE, Kane MS, Litovsky SH, Darley-Usmar VM, Young ME, Chatham JC, Zhang J (2021) Mitochondrial Morphology and Mitophagy in Heart Diseases: Qualitative and Quantitative Analyses Using Transmission Electron Microscopy. Frontiers in Aging 2

Conway AE, Van Nostrand EL, Pratt GA, Aigner S, Wilbert ML, Sundararaman B, Freese P, Lambert NJ, Sathe S, Liang TY et al (2016) Enhanced CLIP Uncovers IMP Protein-RNA Targets in Human Pluripotent Stem Cells Important for Cell Adhesion and Survival. Cell Rep 15: 666–679

Cox J, Mann M (2008) MaxQuant enables high peptide identification rates, individualized p.p.b.-range mass accuracies and proteome-wide protein quantification. Nat Biotechnol 26: 1367–1372

Cox J, Neuhauser N, Michalski A, Scheltema RA, Olsen JV, Mann M (2011) Andromeda: a peptide search engine integrated into the MaxQuant environment. J Proteome Res 10: 1794–1805

Dai N, Christiansen J, Nielsen FC, Avruch J (2013) mTOR complex 2 phosphorylates IMP1 cotranslationally to promote IGF2 production and the proliferation of mouse embryonic fibroblasts. Genes & development 27: 301–312

Dai N, Zhao L, Wrighting D, Kramer D, Majithia A, Wang Y, Cracan V, Borges-Rivera D, Mootha VK, Nahrendorf M et al (2015) IGF2BP2/IMP2-Deficient mice resist obesity through enhanced translation of Ucp1 mRNA and Other mRNAs encoding mitochondrial proteins. Cell metabolism 21: 609–621

deAlmeida AC, van Oort RJ, Wehrens XH (2010) Transverse aortic constriction in mice. J Vis Exp

Degrauwe N, Suva ML, Janiszewska M, Riggi N, Stamenkovic I (2016) IMPs: an RNA-binding protein family that provides a link between stem cell maintenance in normal development and cancer. Genes & development 30: 2459–2474

Dorn LE, Lasman L, Chen J, Xu X, Hund TJ, Medvedovic M, Hanna JH, van Berlo JH, Accornero F (2019) The N(6)-Methyladenosine mRNA Methylase METTL3 Controls Cardiac Homeostasis and Hypertrophy. Circulation 139: 533–545

Duran J, Martinez A, Adler E (2019) Cardiovascular Manifestations of Mitochondrial Disease. Biology (Basel) 8 El-Hattab AW, Scaglia F (2016) Mitochondrial Cardiomyopathies. Front Cardiovasc Med 3: 25

Elachouri G, Vidoni S, Zanna C, Pattyn A, Boukhaddaoui H, Gaget K, Yu-Wai-Man P, Gasparre G, Sarzi E, Delettre C et al (2011) OPA1 links human mitochondrial genome maintenance to mtDNA replication and distribution. Genome Res 21: 12–20

Galindo CL, Skinner MA, Errami M, Olson LD, Watson DA, Li J, McCormick JF, McIver LJ, Kumar NM, Pham TQ et al (2009) Transcriptional profile of isoproterenol-induced cardiomyopathy and comparison to exercise-induced cardiac hypertrophy and human cardiac failure. BMC Physiol 9: 23

Gao C, Wang Y (2020) mRNA Metabolism in Cardiac Development and Disease: Life After Transcription. Physiol Rev 100: 673–694

Huang H, Weng H, Sun W, Qin X, Shi H, Wu H, Zhao BS, Mesquita A, Liu C, Yuan CL et al (2018) Recognition of RNA N(6)-methyladenosine by IGF2BP proteins enhances mRNA stability and translation. Nature cell biology 20: 285–295

Ishikawa TO, Tamai Y, Li Q, Oshima M, Taketo MM (2003) Requirement for tumor suppressor Apc in the morphogenesis of anterior and ventral mouse embryo. Developmental biology 253: 230–246

Janiszewska M, Suva ML, Riggi N, Houtkooper RH, Auwerx J, Clement-Schatlo V, Radovanovic I, Rheinbay E, Provero P, Stamenkovic I (2012) Imp2 controls oxidative phosphorylation and is crucial for preserving glioblastoma cancer stem cells. Genes & development 26: 1926–1944

Kalous A, Stake JI, Yisraeli JK, Holt CE (2014) RNA-binding protein Vg1RBP regulates terminal arbor formation but not long-range axon navigation in the developing visual system. Developmental neurobiology 74: 303–318

Koitabashi N, Bedja D, Zaiman AL, Pinto YM, Zhang M, Gabrielson KL, Takimoto E, Kass DA (2009) Avoidance of transient cardiomyopathy in cardiomyocyte-targeted tamoxifen-induced MerCreMer gene deletion models. Circ Res 105: 12–15

Kolk MV, Meyberg D, Deuse T, Tang-Quan KR, Robbins RC, Reichenspurner H, Schrepfer S (2009) LAD-ligation: a murine model of myocardial infarction. J Vis Exp

Ladha FA, Thakar K, Pettinato AM, Legere N, Ghahremani S, Cohn R, Romano R, Meredith E, Chen YS, Hinson JT (2021) Actinin BioID reveals sarcomere crosstalk with oxidative metabolism through interactions with IGF2BP2. Cell Rep 36: 109512

Laggai S, Kessler SM, Boettcher S, Lebrun V, Gemperlein K, Lederer E, Leclercq IA, Mueller R, Hartmann RW, Haybaeck J et al (2014) The IGF2 mRNA binding protein p62/IGF2BP2-2 induces fatty acid elongation as a critical feature of steatosis. Journal of Lipid Research 55: 1087–1097

Lehnart SE, Wehrens XHT (2022) The role of junctophilin proteins in cellular function. Physiol Rev 102: 1211–1261

Li A, Gao M, Jiang W, Qin Y, Gong G (2020) Mitochondrial Dynamics in Adult Cardiomyocytes and Heart Diseases. Front Cell Dev Biol 8: 584800

Liao Y, Castello A, Fischer B, Leicht S, Foehr S, Frese CK, Ragan C, Kurscheid S, Pagler E, Yang H et al (2016) The Cardiomyocyte RNA-Binding Proteome: Links to Intermediary Metabolism and Heart Disease. Cell Rep 16: 1456–1469

Maggi L, Mavroidis M, Psarras S, Capetanaki Y, Lattanzi G (2021) Skeletal and Cardiac Muscle Disorders Caused by Mutations in Genes Encoding Intermediate Filament Proteins. International Journal of Molecular Sciences 22: 4256

Mathiyalagan P, Adamiak M, Mayourian J, Sassi Y, Liang Y, Agarwal N, Jha D, Zhang S, Kohlbrenner E, Chepurko E et al (2019) FTO-Dependent N(6)-Methyladenosine Regulates Cardiac Function During Remodeling and Repair. Circulation 139: 518–532

May D, Gilon D, Djonov V, Itin A, Lazarus A, Gordon O, Rosenberger C, Keshet E (2008) Transgenic system for conditional induction and rescue of chronic myocardial hibernation provides insights into genomic programs of hibernation. Proc Natl Acad Sci U S A 105: 282–287

McNally EM, Mestroni L (2017) Dilated Cardiomyopathy: Genetic Determinants and Mechanisms. Circ Res 121: 731–748

Orsucci D, Caldarazzo Ienco E, Rossi A, Siciliano G, Mancuso M (2021) Mitochondrial Syndromes Revisited. J Clin Med 10

Pankuweit S, Richter A (2015) Mitochondrial disorders with cardiac dysfunction: an under-reported aetiology with phenotypic heterogeneity. Eur Heart J 36: 2894–2897

Perez-Riverol Y, Bai J, Bandla C, Garcia-Seisdedos D, Hewapathirana S, Kamatchinathan S, Kundu DJ, Prakash A, Frericks-Zipper A, Eisenacher M et al (2022) The PRIDE database resources in 2022: a hub for mass spectrometry-based proteomics evidences. Nucleic Acids Res 50: D543–D552

Pokhrel N, Genin O, Sela-Donenfeld D, Cinnamon Y (2022) HREM, RNAseq and Cell Cycle Analyses Reveal the Role of the G2/M-Regulatory Protein, WEE1, on the Survivability of Chicken Embryos during Diapause. Biomedicines 10

Puspoki Z, Storath M, Sage D, Unser M (2016) Transforms and Operators for Directional Bioimage Analysis: A Survey. Adv Anat Embryol Cell Biol 219: 69–93

Qian B, Wang P, Zhang D, Wu L (2021) m6A modification promotes miR-133a repression during cardiac development and hypertrophy via IGF2BP2. Cell Death Discov 7: 157

Rockman HA, Ross RS, Harris AN, Knowlton KU, Steinhelper ME, Field LJ, Ross J, Jr., Chien KR (1991) Segregation of atrial-specific and inducible expression of an atrial natriuretic factor transgene in an in vivo murine model of cardiac hypertrophy. Proc Natl Acad Sci U S A 88: 8277–8281

Rosenbaum AN, Agre KE, Pereira NL (2020) Genetics of dilated cardiomyopathy: practical implications for heart failure management. Nature reviews Cardiology 17: 286–297

Rosenfeld YB, Krumbein M, Yeffet A, Schiffmann N, Mishalian I, Pikarsky E, Oberman F, Fridlender Z, Yisraeli JK (2019) VICKZ1 enhances tumor progression and metastasis in lung adenocarcinomas in mice. Oncogene 38: 4169–4181

Scheltema RA, Hauschild JP, Lange O, Hornburg D, Denisov E, Damoc E, Kuehn A, Makarov A, Mann M (2014) The Q Exactive HF, a Benchtop mass spectrometer with a pre-filter, high-performance quadrupole and an ultra-high-field Orbitrap analyzer. Mol Cell Proteomics 13: 3698–3708

Stege NM, de Boer RA, van den Berg MP, Silljé HHW (2021) The Time Has Come to Explore Plasma Biomarkers in Genetic Cardiomyopathies. International Journal of Molecular Sciences 22: 2955

Tarnavski O, McMullen JR, Schinke M, Nie Q, Kong S, Izumo S (2004) Mouse cardiac surgery: comprehensive techniques for the generation of mouse models of human diseases and their application for genomic studies. Physiol Genomics 16: 349–360

Tybl E, Shi F-D, Kessler SM, Tierling S, Walter J, Bohle RM, Wieland S, Zhang J, Tan EM, Kiemer AK (2011) Overexpression of the IGF2-mRNA binding protein p62 in transgenic mice induces a steatotic phenotype. J Hepatol 54: 994–1001

Wai T, Garcia-Prieto J, Baker MJ, Merkwirth C, Benit P, Rustin P, Ruperez FJ, Barbas C, Ibanez B, Langer T (2015) Imbalanced OPA1 processing and mitochondrial fragmentation cause heart failure in mice. Science 350: aad0116

Wessel D, Flugge UI (1984) A method for the quantitative recovery of protein in dilute solution in the presence of detergents and lipids. Anal Biochem 138: 141–143

Wu W, Ziemann M, Huynh K, She G, Pang ZD, Zhang Y, Duong T, Kiriazis H, Pu TT, Bai RY et al (2021) Activation of Hippo signaling pathway mediates mitochondria dysfunction and dilated cardiomyopathy in mice. Theranostics 11: 8993–9008

Yisraeli JK (2005) VICKZ proteins: a multi-talented family of regulatory RNA-binding proteins. Biol Cell 97: 87–96

Zhou B, Tian R (2018) Mitochondrial dysfunction in pathophysiology of heart failure. The Journal of clinical investigation 128: 3716–3726

