## Supplemental figure file for "IGF2BP2 is Induced by Stress in the Heart and Mediates Dilated Cardiomyopathy"

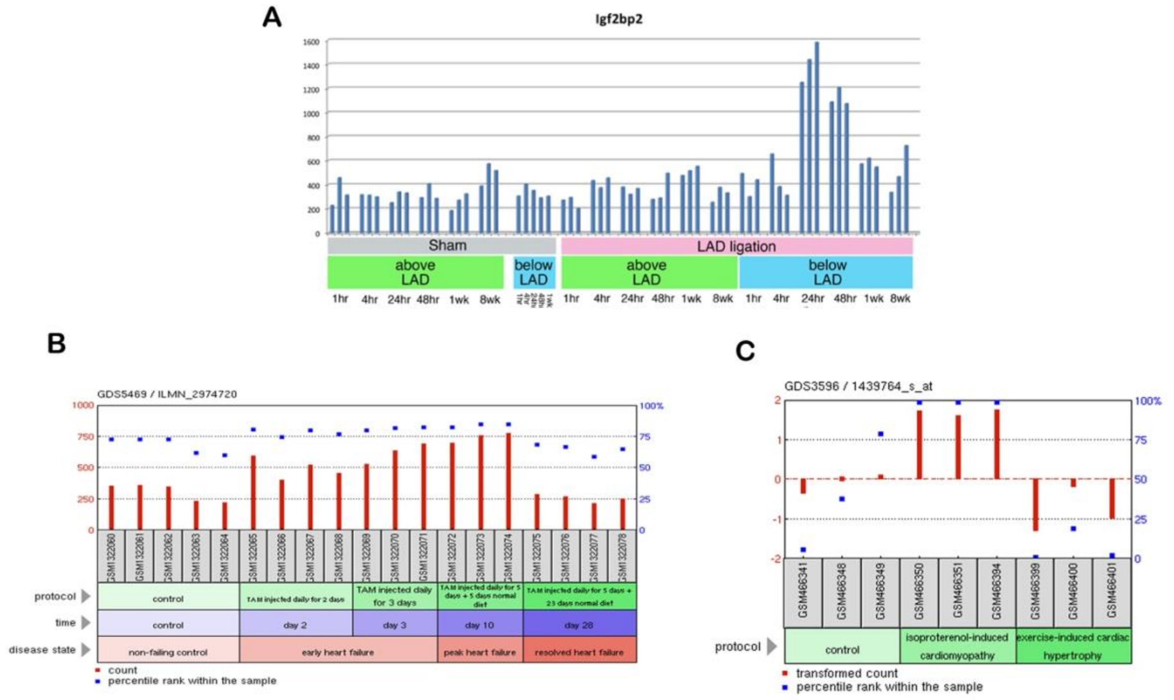


Figure S1. <u>Cardiac stress models and IGF2BP2 expression</u>
(A) IGF2BP2 mRNA expression was assayed, from 1 hour to 8 weeks, in Sham-operated and LAD ligated mice at levels above (unaffected area) and below (affected area) of the LAD (GEO data set GDS488). (B) Microarray data of IGF2BP2 expression from early (2,3 days), late (10 days), and resolved (28 days) heart failure in a MerCreMer mouse model (GEO data set GDS5469). IGF2BP2 mRNA expression is upregulated already at early stages and then down regulated upon recovery. (C) Microarray data of IGF2BP2 expression in isoproterenol-induced cardiomyopathy compared to exercised induced cardiac hypertrophy (GEO data set GDS3596). IGF2BP2 mRNA expression is upregulated in isoproterenol-induced cardiomyopathy and unchanged in exercise induced cardiac hypertrophy.

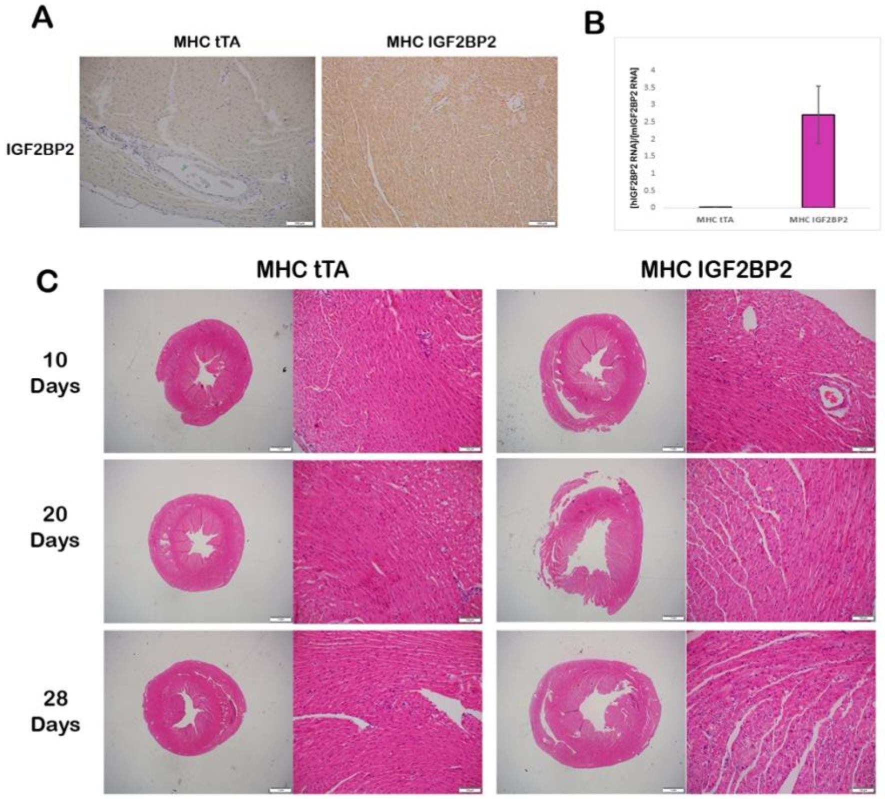


Figure S2. <u>Expression of IGF2BP paralogs</u>
(A) Relative IGF2BP2 RNA expression was measured by RT-PCR in normal hearts at post-natal day 1 (P1), post-natal day 8 (P8), 3 weeks (3W) and 6 weeks (6W) of age.  Each bar represents three biological repeats ± SEM). (B) Western blot of IGF2BPs at the indicated time points. α/β Tubulin was used as a loading control. On the right, quantification of western results. Two biological repeats are shown for each time point ± SEM. (C) Hearts from MHC-tTA and MHC-tTA/IGF2BP2 mice were stained for IGF2BP2 expression with an anti-IGF2BP2 antibody. Scale bar is 100µm. (D)  The level of transgene expression (normalized to endogenous expression; hIGF2BP2/mIGF2BP2) in primary cardiomyocytes was measured by RT-PCR 5 days after transgene induction (withdrawal from tetracycline).  Each bar represents 4 biological repeats ± SEM. (E) H&E staining of MHC-tTA and MHC-tTA/IGF2BP2 hearts at the time points indicated after the beginning of transgene induction. Scale bar is 1mm and 100µm in the higher magnification (right panels).

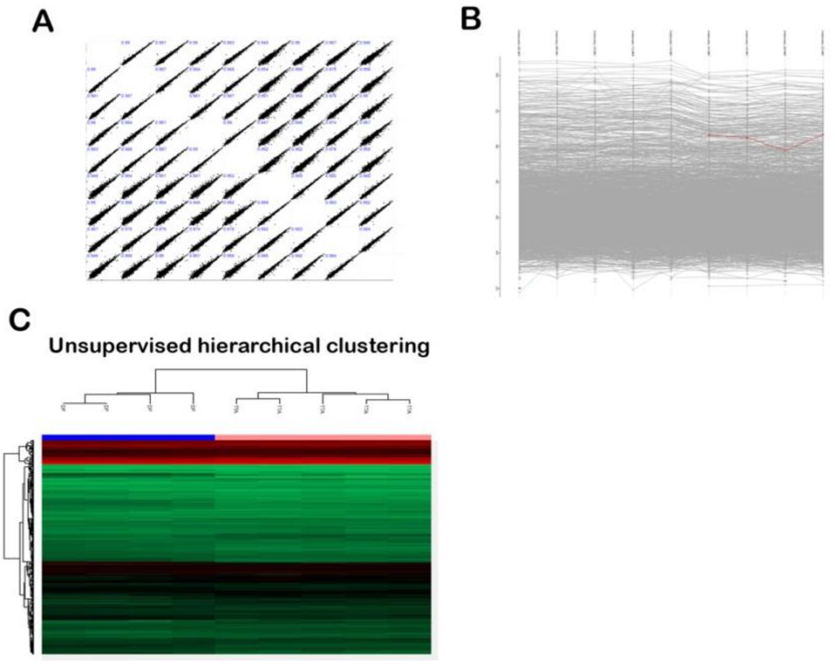


Figure S3. Data from Mass Spectrometry analysis
(A) A multi scatter plot with Pearson correlations of all 9 samples described in figure 5. (B) IGF2BP2 profile plot in all 9 samples. (C) Unsupervised hierarchical clustering of all 9 samples without Z score. Orange header, MHC tTA samples; Blue header, MHC IGF2BP2 samples.

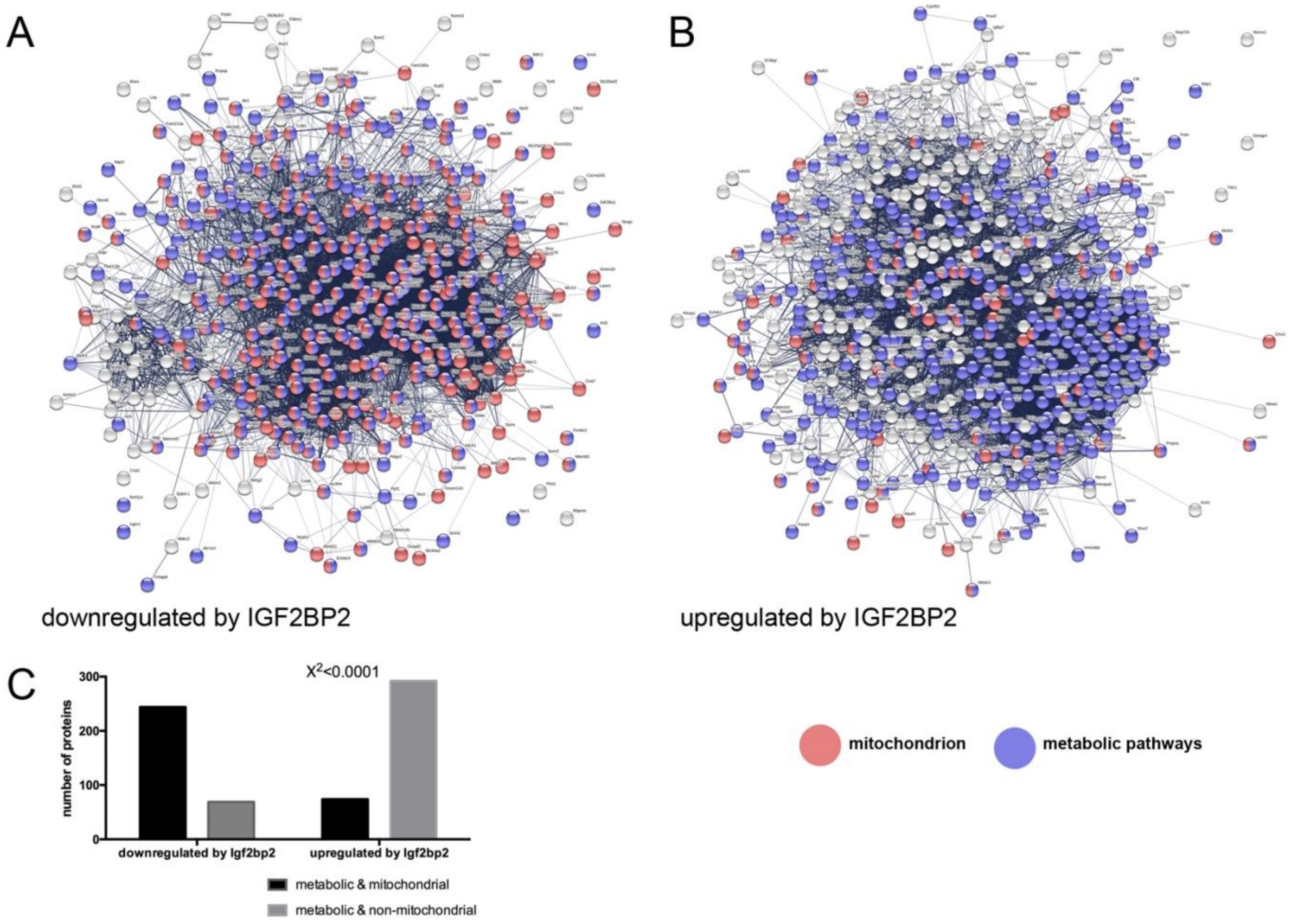


Figure S4. Protein-protein interaction network comparison of proteins down- or up-regulated by IGF2BP2 overexpression.

Proteins significantly down- (A) or up- (B) regulated (FDR=0.05; S=0.1) in the MS analysis (Supplemental File 1) were displayed in a Protein-Protein interaction analysis using the STRING website. Genes associated with the Mitochondrion Cellular Components gene set (GO:0005739) were colored pink and those associated with Metabolic Pathways Biological Processes (GO:0008152) were colored blue. A highly significant overlap ($\chi^2<0.0001$) of metabolic and mitochondrial proteins is present in the downregulated as opposed to upregulated proteins (C).

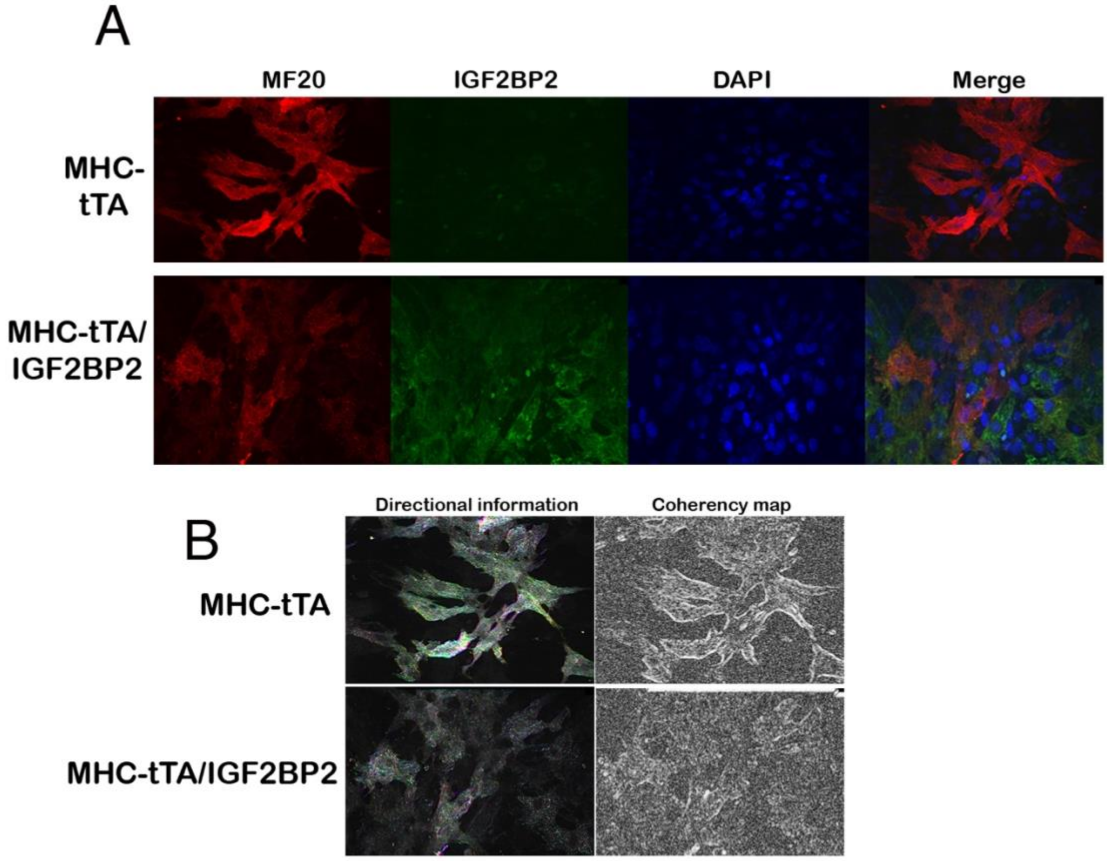


Figure S5. IGF2BP2 expression in primary cardiomyocytes affects sarcomere structure
(A) Primary cardiomyocytes isolated from the hearts of either MHC-tTA or MHC-tTA/IGF2BP2 mice were grown for 7 days in culture in the absence of tetracycline (to activate the transgene) and then stained for MF20, IGF2BP2 and DAPI. (B) Analysis using ImageJ OrientationJ plugin of directionality (left) and coherency (right) of the pixels of MF20 staining (346 MHC-tTA cells and 216 MHC-tTA/IGF2BP2 cells were counted).

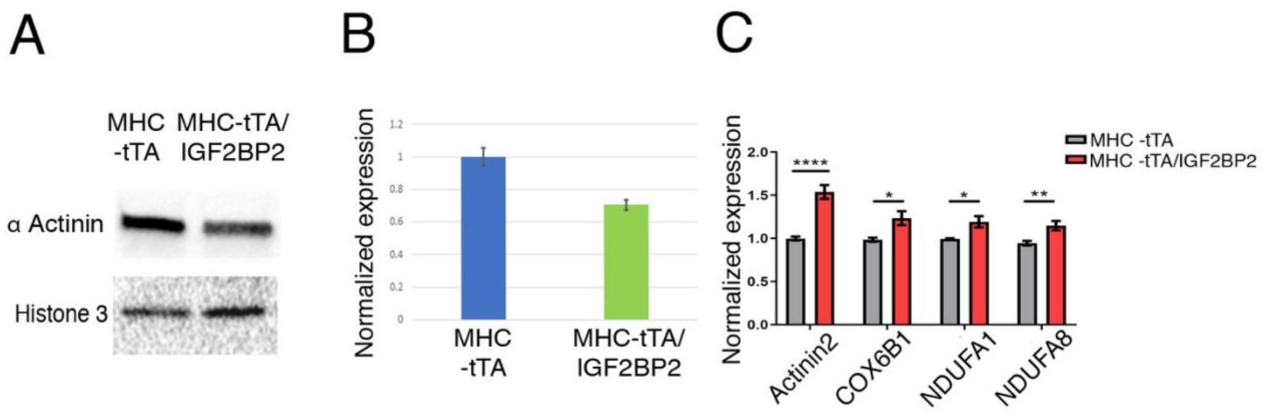


Figure S6. Protein and mRNA expression induced by IGF2BP2 expression in the heart.
(A) Western blot of proteins extracted from MHC-tTA and MHC-tTA/IGF2BP2 mice expressing the transgene for 15 days in the heart, using a pan α Actinin antibody. (B) Quantification of (A) based on two biological repeats ±SEM. (C) RT- PCR of RNA isolated from PCMs 5 days after IGF2BP2 expression was activated. The results represent the average of 3 biological repeats, with each repeat evaluated in duplicate. Statistical significance was determined by two tailed Student's t-test: * $p<0.05$, ** $p<0.01$, ****$p<0.0001$
